# Predicting GD2 expression across cancer types by the integration of pathway topology and transcriptome data

**DOI:** 10.1101/2025.05.21.655303

**Authors:** Arsenij Ustjanzew, Federico Marini, Saskia Wagner, Arthur Wingerter, Roger Sandhoff, Jörg Faber, Claudia Paret

## Abstract

The disialoganglioside GD2 is a key cancer therapy target due to its overexpression in several cancer types and limited expression in normal tissues. We developed a computational framework integrating reaction activity scores derived from transcriptomic data with glycosphingolipid biosynthesis pathway to predict GD2 expression. We labeled specific reactions as GD2-promoting or -mitigating, and used their cumulative activity as features to distinguish neuroblastoma from normal tissue. Predicted GD2 scores were validated by comparing them with literature-reported values and by assessing GD2 expression through flow cytometry in clear cell sarcoma of the kidney, which express GD2 at high level according to the GD2 scores. GD2 expression heterogeneity across cancer subtypes indicated patient subgroups potentially suitable for targeted therapy, and a role of *B4GALNT1* amplification as a GD2 promoting factor. We offer our approach via the R package GD2Viz including also an interactive Shiny application and enable an enhanced processing of custom datasets.

## 1. Introduction

Gangliosides are a group of sialylated glycosphingolipids (GSL) involved in differentiation, cell signaling, adhesion, and neuronal development ^1–3^. Structurally, they are composed of a ceramide tail, attached to an oligosaccharide moiety and one or more sialic acid residues ^4^. The ganglioside biosynthesis pathway is a stepwise process that involves multiple enzymes, beginning with the formation of glycosylceramide from ceramide by the enzyme encoded from *UGCG* and continuing with the sequential addition of saccharides through a variety of glycosyl-and sialyltransferases. The initial enzymes (encoded by *ST3GAL3/5* and *ST8SIA1*) in this pathway demonstrate high substrate specificity, directing the formation of distinct ganglioside series (0-, a-, b-, and c-series), while downstream enzymes, such as those encoded by *B4GALNT1* and *B3GALT4*, exhibit broader substrate specificity and elongate all four series ^4,5^. The disialoganglioside GD2, which is synthesized from its precursor GD3 by the transferase encoded by *B4GALNT1*, is of particular importance due to its functional involvement in developmental processes and its pathophysiological roles in cancer diseases^6^_._

The expression of GD2 is limited in normal human tissues, particularly described in the central nervous system and peripheral sensory nerve fibers ^6^. Conversely, GD2 is overexpressed in several tumor types, including neuroblastoma (NB), melanoma, small-cell lung cancer, and various sarcomas ^6,7^. This makes GD2 a promising target for therapeutic interventions, such as monoclonal antibodies and CAR-T cell therapies ^8–10^. Among the aforementioned cancers, neuroblastoma exhibits particularly high levels of GD2 expression and antibodies against GD2 are already included in the treatment protocol of high-risk NB ^11^. Currently, 23 clinical studies listed on ClinicalTrials.gov are recruiting patients from various tumor types for GD2-based therapies (search term: https://clinicaltrials.gov/search?cond=GD2&aggFilters=status:rec, received on 2025-02-06). However, GD2 expression varies both across and within different tumor types ^12^. Therefore, assessing GD2 expression is crucial for optimizing patient enrollment and should be performed prior to administering anti-GD2 therapies.

At present, the measurement of GD2 expression in tumors can be conducted through techniques such as immunohistochemistry (IHC), thin-layer chromatography (TLC), mass spectrometry, and flow cytometry ^7^. The use of IHC on formalin-fixed, paraffin-embedded samples is limited by the potential for lipids to be washed out by the solvent used in tissue processing ^13^. Other methods, such as TLC and liquid chromatography-coupled tandem mass spectrometry, are more specific but require specialized equipment and are costly and labor-intensive ^12^. Flow cytometry requires a cell suspension which is generally not available in routine diagnostics. Consequently, these methodologies are not readily available for routine clinical applications, particularly in resource-limited settings; moreover, no diagnostic assays are available on the market to detect GD2.

RNA sequencing (RNA-seq) has emerged as a powerful tool in the field of clinical oncology, offering a comprehensive view of gene expression. In molecular tumor boards, where personalized cancer treatments are discussed, RNA-seq data is employed to identify fusions and overexpressed molecular targets ^14,15^. GD2 expression is not typically analyzed in these settings due to the complexity of lipid detection. RNA-seq could present an opportunity to assess ganglioside-related gene activity, offering detailed gene expression profiles that could potentially be used to infer GD2 levels. However, translating transcriptomic data into accurate ganglioside phenotype predictions remains a challenge, underscoring the need for methods and models that bridge the gap between gene expression and metabolic profiles.

Previous methods for predicting GD2 expression have primarily relied on individual genes or small gene signatures ^16,17^. For example, *Sorokov et al.* 2020 proposed a two-gene signature comprising *ST8SIA1* and *B4GALNT1*, aiming to correlate their expression with a GD2-positive phenotype ^18^. Similarly, *Sha et al.* 2021 developed a glycosyltransferase score based on neuroblastoma datasets, which correlated with GD2 positivity ^19^. However, these models are limited by their narrow focus on single genes or cancer types, neglecting the broader metabolic pathway and enzyme-substrate specificities involved in ganglioside metabolism.

In this study, we hypothesized that the integration of gene expression and ganglioside biosynthesis pathway information, namely the specific labeling of reactions that promote GD2 (participating in the sequential biosynthesis of GD2) and mitigate GD2 (downstream metabolism of GD2 or entry into neighboring ganglioside series), may serve as an effective predictor of the GD2-positive phenotype. We used publicly available RNA-seq datasets from NB tumor and normal tissue samples to compute Reaction Activity Scores (RAS). To address the ambiguity regarding the substrate specificity of certain enzymes in ganglioside metabolism, we adjusted the RAS values with transition probability of the pathway topology^20^. The adjusted RAS values of GD2-promoting and -mitigating reactions were then utilized to train a Support Vector Machine (SVM) model. We evaluated different kernels, and validated the GD2 score against additional public datasets and discussed it in context of previous literature knowledge regarding GD2 expression in different tumor entities. Finally, we developed the R package GD2Viz, which is available at https://github.com/arsenij-ust/GD2Viz, to provide scientists and clinicians easier access to our methodology. The package contains R functions that facilitate the prediction of the GD2 score, encapsulated in an interactive Shiny app that enables users to efficiently process their datasets and explore them in the context of GD2 prediction.

In conclusion, we have constructed a machine learning model that can be utilized as a predictor for GD2 molecular phenotypes in diverse tumor entities, along with the GD2Viz tool, which can assist in diagnostic decisions and provide deeper insights into the role of GD2 in cancer biology.

## 2. Materials and Methods

### 2.1 Data Acquisition and Preprocessing

For the training dataset, the gene expression data (RSEM expected counts) and clinical information of the combined TCGA-TARGET-GTEx cohort (n=19,109) were obtained from the UCSC Xena database (http://xena.ucsc.edu/) ^21^ as log2(expected count + 1) (data was retrieved on 2022-10-06). This cohort is the result of the UCSC Toil RNA-seq recompute compendium ^22^. The gene counts were subsequently transformed back to integer counts. The dataset was subset to neuroblastoma samples from the Therapeutically Applicable Research to Generate Effective Treatments (TARGET) project ^23^ and normal tissue samples from the Genotype-Tissue Expression (GTEx) project ^24^. Additional clinical information was derived from the Genomic Data Commons (GDC) Data Portal and merged according to the case submitter IDs ^25^. The removal of cell lines and ganglioneuroblastoma samples resulted in a total of 7,548 samples, of which 136 were identified as NB. The Ensembl IDs were annotated to gene symbols using the R package org.Hs.eg.db (version 3.14.0) ^26^, resulting in 34,281 genes. The gene counts were normalized using the median of ratios method provided by the DESeq2 R package (version 1.34.0) ^27^. Finally, a pseudocount was added and the counts were log10 transformed.

GD2 scores were computed for the following publicly available datasets: *TCGA*: As stated above, the TCGA-TARGET-GTEx cohort was subset to TCGA project samples (n=10,529). TCGA molecular subtypes were retrieved using the TCGAbiolinks R package (version 2.22.4) ^28^. Additionally, molecular subtype information for TCGA breast cancer (BRCA) was retrieved as published in *Lehmann et al.* ^29^. For the cancer subtype analysis, the TCGA dataset was divided into subprojects. The Ensembl IDs were annotated to gene symbols. The copy number alteration (CNA) data of TCGA BRCA, glioblastoma multiorme (GBM), low-grade glioma (LGG), lung adenocarcinoma (LUAD), and sarcoma (SARC) was separately retrieved from the official cBioPortal for Cancer Genomics portal (https://www.cbioportal.org/) (data was retrieved on 2025-01-21).

*TARGET*: As stated above, the TCGA-TARGET-GTEx cohort was subset to TARGET project samples (n=734). The Ensembl IDs were annotated to gene symbols. Tumor entities containing less than five samples per category were excluded. ^23^

*GTEx*: Similarly, the TCGA-TARGET-GTEx cohort was reduced to GTEx project samples. The Ensembl IDs were annotated to gene symbols. Tissue types containing less than five samples per category were excluded. Cell line samples were also excluded from the heatmap visualization resulting in a total of 7,845 samples. ^24^

*St. Jude Cloud*: RNA-seq profiles and corresponding annotation of 2,853 samples were obtained from the St. Jude Cloud (https://stjude.cloud) (data was retrieved on 2023-05-12). We excluded genes with less than 10 read counts in total of all samples. The batch effect caused by different library preparation protocols was removed using the ComBat_seq function from the sva R package (version 3.42.0). Tumor entities containing less than five samples per category were excluded.

*CBTTC*: The gene expression data and clinical information of the Pediatric Brain Tumor Atlas: Children’s Brain Tumor Tissue Consortium CBTTC cohort (n=970) were obtained from the UCSC Xena database as log2(expected_count + 1) (data was retrieved on 2024-06-28). The gene counts were subsequently transformed back to integer counts (https://cbttc.org). ^30^

*GSE117446*: Raw count matrix and genotype of 29 midline high-grade gliomas were retrieved from Gene Expression Omnibus (GEO accession: GSE117446). The dataset contained 16 H3K27M mutated and 13 H3 wildtype samples (data was retrieved on 2023-09-27). The dataset included the Ensembl gene IDs and gene symbols. The gene symbols were used in further analysis. ^31^

*GSE147635*: Expression data and clinical information were obtained from Gene Expression Omnibus (GEO accession: GSE147635). The dataset contained 6 ganglioneuroma (GN) and 15 neuroblastoma (NB) samples ^32^ (data was retrieved on 2023-02-13). The processing of raw RNA-seq data for this dataset was described in our previous work (*Ustjanzew et al.* 2024 ^20^).

*GSE180514*: Raw RNA-seq counts and GD2 status of 8 neuroblastoma Kelly cells were retrieved from Gene Expression Omnibus (GEO accession: GSE180514). The dataset contained 4 GD2-low and 4 GD2-high samples, where GD2 level was identified by staining with GD2-APC antibody and FACS-sorted into the respective category. The Homo sapiens Annotation Release 109.20190905 was used for gene annotation (data was retrieved on 2023-11-21). ^16^

### 2.2 Workflow Overview

Figure 1 provides a schematic overview of the computational pipeline for model development and GD2 score prediction. The workflow starts with the construction of a GSL metabolic network by merging four Kyoto Encyclopedia of Genes and Genomes (KEGG) pathways related to the Sphingolipid metabolism. In this metabolite-centric network, metabolites are represented as nodes, while biochemical reactions define edges. Next, Reaction Activity Scores (RAS) are derived from RNA-seq data to quantify reaction activities across individual samples. These scores are subsequently used to assign weights to the metabolic network edges, enabling sample-specific reaction profiling. To refine reaction-level resolution, we incorporated network topology by integrating transition probability between nodes with RAS values. This approach facilitates the disambiguation of enzymatic substrate specificity. In this study, we systematically assessed the impact of raw RAS values and topology-adjusted reactions on the computed GD2 score. Reactions were categorized based on their influence on GD2 levels, with mitigating (GD2r-) and promoting (GD2r+) reactions identified, according to their position in the metabolic pathway.

**Figure 1.**
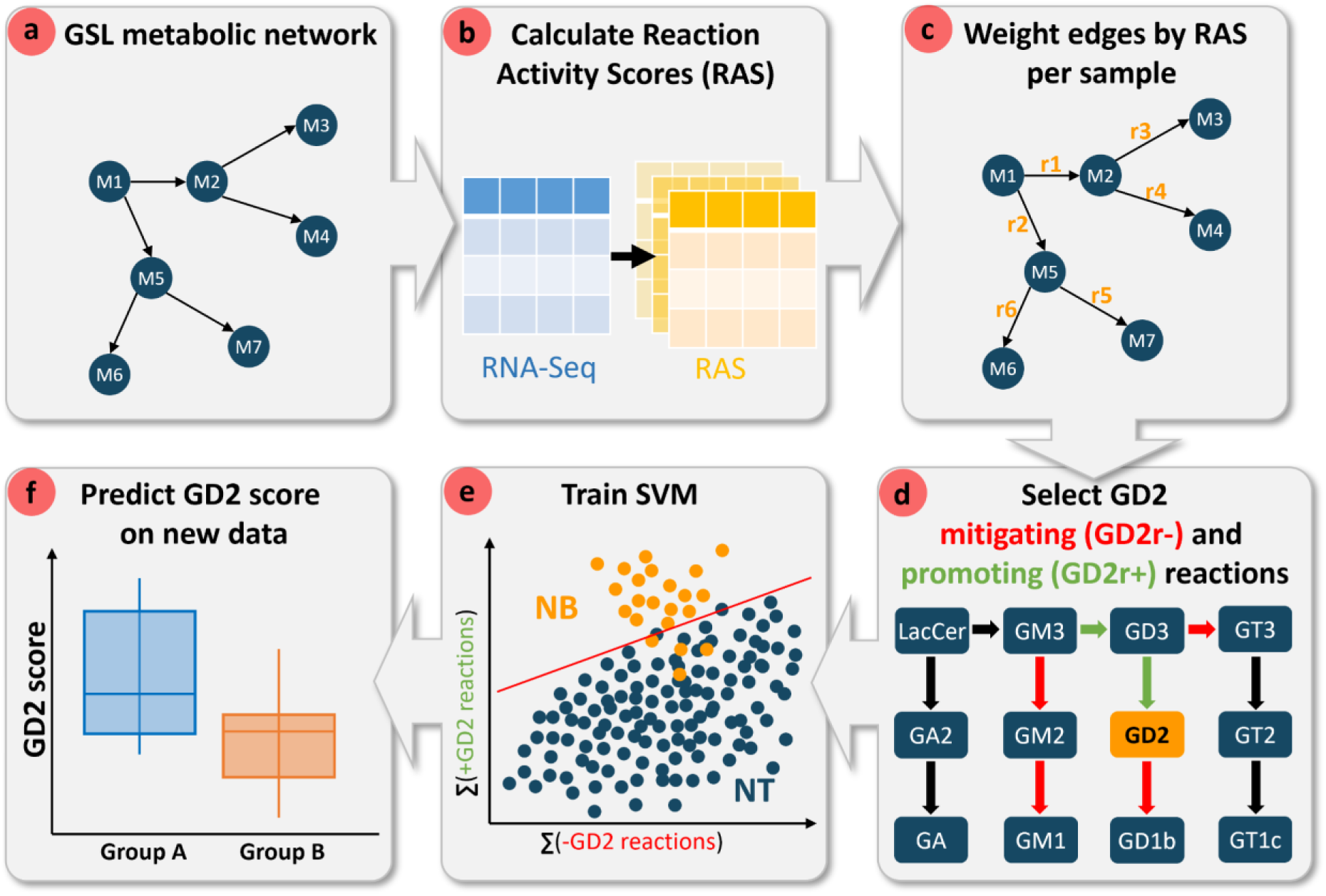
W**o**rkflow **diagram of the computational pipeline for predicting GD2 scores using transcriptome data and metabolic network topology.** a) A glycosphingolipid (GSL) metabolic network is constructed, where nodes represent metabolites and edges denote biochemical reactions. b) Reaction Activity Scores (RAS) are computed from RNA-Seq data to quantify reaction activities across samples. c) The metabolic network is weighted based on RAS values for each sample. d) Reactions are categorized as either mitigating (GD2r-, red) or promoting (GD2r+, green) based on their position in the metabolic pathway in respect to GD2. e) A support vector machine (SVM) classifier is trained using the cumulative activity of GD2r- and GD2r+ reactions to distinguish neuroblastoma (NB) from normal tissue (NT) samples. f) The trained model is then applied to predict GD2 scores in independent datasets, facilitating group comparisons.

A SVM classifier was trained using the cumulative activity of the +GD2 and -GD2 reactions to distinguish NB from normal tissue (NT) samples. Various combinations of +GD2 and -GD2 reactions, as well as different SVM kernels, were evaluated to optimize model performance and explainability. Finally, the trained model was applied to predict GD2 scores in independent datasets, which were used for subsequent validation based on expected biological outcomes and existing literature. The GD2 score enables comparative analyses across different biological groups, such as cancer subtypes, and may further serve as a basis for the identification of possible biomarkers and cancer-associated genes linked to GD2 expression. The workflow was executed fully in R (version 4.1.3).

### 2.3 Graph Construction

A metabolite-centric graph was constructed using the R packages *NetPathMiner* (v1.30.0) and *igraph* (v1.4.2) using data from the Kyoto Encyclopedia of Genes and Genomes (KEGG). Nodes represent metabolites, and edges connect metabolites involved in the same reactions, with edge metadata specifying the responsible enzymes. The graph includes four KEGG pathways (hsa00600, hsa00601, hsa00603, hsa00604) covering sphingolipid and glycosphingolipid metabolism, including the lacto-, neolacto-, globo-, and ganglio-series. Degradation reactions R06010 (GM1 to GM2) and R06004 (GM2 to GM3) were excluded as they are not competing enzymatic reactions of the biosynthesis resulting in a metabolic graph containing in total 104 nodes and 116 edges. KEGG data were retrieved on 2022-07-07.

### 2.4 Weighting of the graph

In the following section, we provide a brief overview of the methodology introduced in *Ustjanzew et al.* 2024 to integrate the pathway topology with transcriptomic data ^20^. For each sample, the metabolic graph was weighted using RAS and integrated with transition probabilities. The weighting of the metabolic graph is initiated by the computation of RAS values based on gene expression levels and gene-protein-reaction (GPR) rules, which serve as a proxy for the activity of enzymatic reactions. In order to account for the lack of enzyme specificity in the ganglioside metabolism pathway, the RAS values were adjusted by a topological metric, namely the RAS-based transition probabilities of the metabolic graph. For a more detailed explanation of the RAS adjustment methods, please refer to *Ustjanzew et al.* 2024.

#### Calculating Reaction Activity Scores

The edges in the metabolic graph were assigned weights based on RAS, a measure of enzymatic activity for each reaction in the graph, as introduced by *Graudenzi et al.* 2018 ^33^. The RAS was calculated based on the decimal normalized gene expression levels from patient samples and the GPR association rules derived from the *Homo sapiens* genome-scale metabolic model (Human1) ^34^. GPRs are logical formulas that describe the enzymes involved in specific reactions and how they relate to one another when multiple enzymes are associated with a common reaction. The minimum gene expression levels were assigned to reactions with an "AND" operator (e.g., multiple subunits), while those with an "OR" operator (e.g., isoenzymes) got the sum of gene expression levels across the involved genes.

#### Adjusting RAS by Transition Probabilities

For each sample we calculated a transition probability (TP) matrix, representing the likelihood of transitioning from one node to the next in a single step in the metabolic graph based on the RAS values of outgoing edges. The TP for an outgoing edge of node x was calculated by dividing the edge’s RAS value by the sum of all outgoing edges for that node. The edge weights were adjusted by multiplying the TPs by the RAS values.

To address the challenge of low enzyme specificity, particularly along the 0-, a-, b-, and c-series within the ganglioside metabolism pathway, we employed an alternative TP matrix to adjust the RAS values. In this alternative adjustment method (hereafter, we will refer to this method as the recursive adjustment), the TPs that were equal to 1 (due to their status as the sole outgoing edges) were assigned the next available upstream TP that was not equal to 1. As with the original TP, the alternative TP matrix was multiplied by the RAS values per sample.

### 2.5 Model Construction & Training

Starting with (un-) adjusted RAS matrices (depending on the RAS adjustment method applied) of the training dataset, where columns correspond to reactions and rows to samples, GD2-promoting reactions (GD2r+), namely R05946, R05940, and GD2-mitigating reactions (GD2r-), namely R05939, R05948, R05947, R05941, were selected based on their position in the gangliosphingolipid metabolism pathway (Figure 2). Hereafter we refer to this selection as *ab initio* reactions. The summed RAS of GD2r+ and GD2r-were used as two predictor features. The data was formatted into a binary classification task to distinguish TARGET NB samples from GTEx samples.

**Figure 2.**
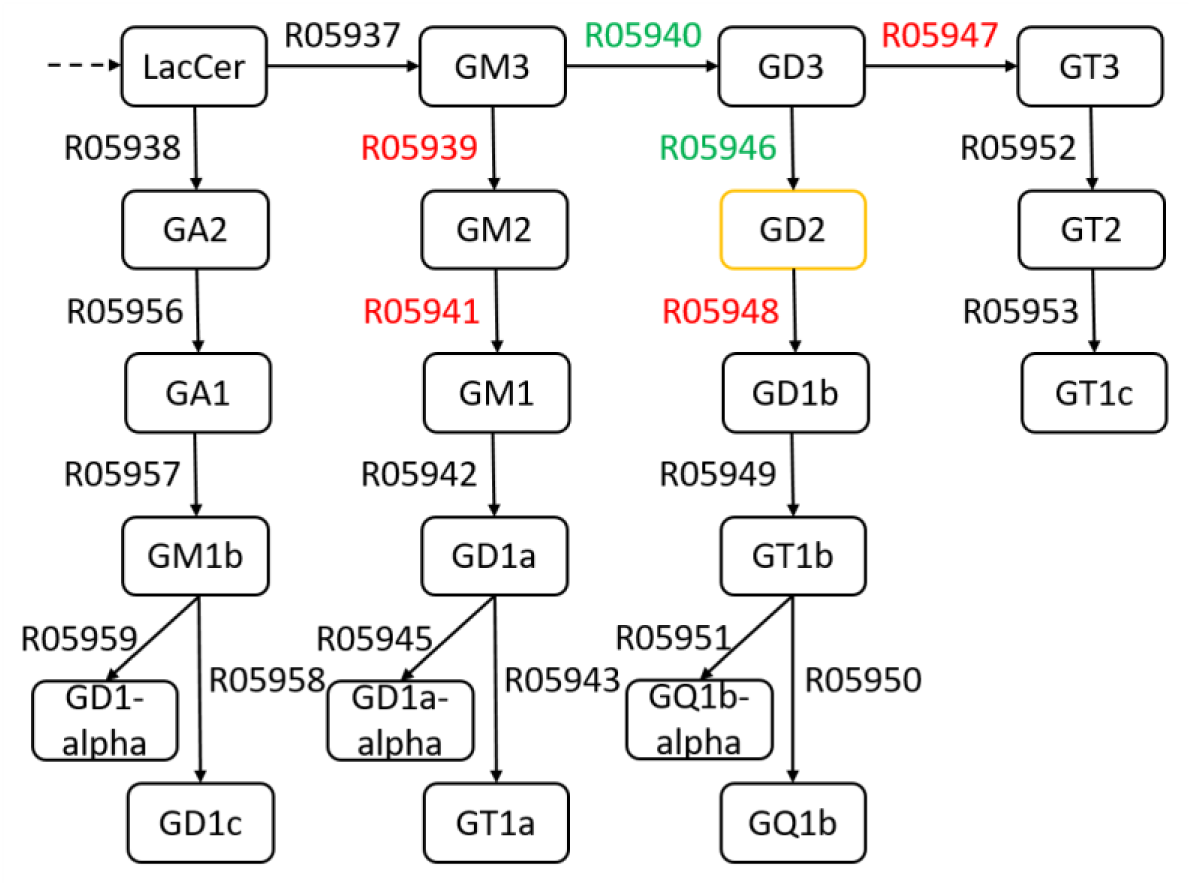
Partial view of the constructed GSL network based on the KEGG-pathway of the glycosphingolipid biosynthesis -ganglio series. *Ab initio*-defined GDr+ IDs are green and GD2r-IDs are red.

For model training, we employed the ksvm function from the kernlab package (version 0.9-32) configuring the SVM with a linear kernel and fitting the model to the classification problem. In the remainder of this work, we refer to the GD2 score based on RAS values as *ras*, based on TP-adjusted RAS values as *rasTP*, and based on recursively adjusted RAS values as *rasTPrec*.

### 2.6 Model evaluation

#### Evaluation of GD2r+/GD2r-

To validate whether the *ab initio*-defined GD2r+ and GD2r-predictive variables contribute meaningfully to the classification task, a permutation test was conducted on the training dataset with 10,000 random permutations of GSL reaction labels (two random reaction labels for GD2r+ and four for GD2r-). The null hypothesis was that the *ab initio*-defined reactions of GD2r+ and GD2r-do not have any true predictive relevance. The binary class predictions (TARGET NB samples labeled as 1 and GTEx samples as 0) yielded empirical p-values for different model quality metrics by the caret package (version 6.0-94). Due to the unbalanced nature of the training dataset, we employed four quality metrics to ascertain whether the model exhibited better performance than that of a classifier based on random reactions: balanced accuracy, precision, F1-score, and recall. p-values were computed as the number of permuted metrics with bigger values compared to the observed metric of the *ab initio* model divided by the total number of permutations.

Additionally, the reactions of GD2r+ and GD2r-were manually altered based on pathway-plausible considerations, resulting in 16 models with varying reaction combinations. These models were subject to equal validation through the aforementioned quality metric calculation.

#### Alternative kernel functions

Further exploration involved training SVM models with alternative kernel functions, such as radial basis, polynomial, hyperbolic tangent, Laplace, Bessel, and ANOVA. Each kernel was evaluated across varying hyperparameters to determine their effect on classification performance. The hyperparameter values are reported in Supplemental file 1.

#### Prediction of GD2 scores

To ensure scale consistency and homoscedasticity between the training and independent datasets in the prediction of GD2 scores, the RNA-seq data from the independent datasets must undergo the same normalization and transformation steps as the training set ^35^. As the DESeq2 median of ratios method was employed for the normalization of the training dataset, the size factors (s*) of the new sets (x*) should be estimated according to the following formula:

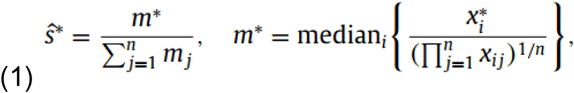

where x denotes a p × n dimensional RNA-seq gene expression data matrix, with p genes and n samples. Let x_ij_ be the elements of RNA sequencing count matrix for i-th gene (i = 1, 2, . . ., p) and j-th sample (j = 1, 2, . . ., n). m* and s* are estimated using the geometric means derived from the training data.

Following normalization and log10-transformation, the RAS values for the GSL metabolism were calculated as previously described using the normalized decimal logarithmic gene expression values. The TP for each sample was then calculated and employed to adjust the RAS accordingly. Subsequently, the *ab initio* defined reactions were aggregated into GD2r+ and GD2r-variables for both the adjusted and unadjusted RAS datasets, enabling the prediction of decision values through the respective SVM model. These decision values provided quantitative estimates of the GD2 score across the test datasets.

### 2.6 GD2 Two-gene Signature

The normalized log10-transformed gene expression of *B4GALNT1* and *ST8SIA1* were summed to calculate the two-gene-signature, as it was defined in the work of *Sorokov et al.* (2020) ^18^.

### 2.7 Identification of CNA associated with the GD2 score

Statistical analyses were conducted to evaluate the relationship between the CNA status of genes and the GD2 score across subgroups of TCGA GBM, TCGA SARC, and TCGA LUAD.

The processed CNA data for the respective TCGA projects were downloaded from the cBioPortal platform (data was retrieved on 2025-01-22). CNA data and gene expression data were integrated by Patient ID and the intersection of common genes (annotated in both as HUGO gene symbols). The CNA status levels -2, -1, 0, 1, and 2 were renamed to homozygous deletion, hemizygous deletion, no change, gain, and high-level amplification. Following filtering criteria were applied to ensure robust statistical analysis: Samples with missing CNA status values were excluded from the analysis. A minimum of 5 samples was required per subgroup-gene-CNA status combination. Genes were included only if they exhibited more than one unique CNA status level per group.

### 2.8 Statistical Tests

Wilcoxon rank-sum tests were performed to assess differences in GD2 scores (based on *ras*, *rasTP*, *rasTPrec*) and two-gene signatures for two groups. Cliffs’s delta was computed with the effsize package (version 0.8.1). For the differences across more than two groups, the Kruskal-Wallis test was used. Dunn’s test was applied for post hoc testing using the rstatix package (version 0.7.2).

Kruskal-Wallis tests were performed to assess differences in GD2 scores across CNA status levels for each gene within each subgroup. Analyses were conducted separately for each subgroup of interest. Eta-squared estimates (η^2^) for the Kruskal-Wallis tests were calculated using the rstatix package (version 0.7.2). To control for the false discovery rate (FDR) in the context of multiple hypothesis testing, p-values were adjusted using the Benjamini-Hochberg method ^36^.

Genes were considered statistically significant if 1) the adjusted p-value was less than 0.05 and 2) the η^2^ exceeded 0.06, indicating a moderate or strong association.

### 2.9 Oncogenic Gene Comparison

Significant genes from each group were cross-referenced with a curated list of oncogenic targets from OncoKB, encompassing 1,172 genes (https://www.oncokb.org/cancer-genes, OncoKB list version: last updated on 2024-12-19 was retrieved on 2025-01-20). Gene overlap was identified to prioritize potential oncogenic drivers associated with the GD2 score.

### 2.10 Cell Culture

Human neuroblastoma cell lines acquired from DMSZ (Braunschweig, Germany) were kept at 37°C and 5% CO₂ in 10 cm² cell culture dishes (SARSTEDT AG & Co. KG, Nümbrecht, Germany). CHP-134 cells were cultivated in RPMI-1640 medium (Sigma Life Science, St. Louis, USA), and SH-SY5Y cells were cultivated in Advanced DMEM (Sigma Life Science, St. Louis, USA). 1% penicillin/streptomycin (Dickinson and Company, Franklin Lakes, USA), 1% L-glutamine (Sigma Life Science, St. Louis, USA), and 10% FCS (Thermo Fisher Scientific GmbH, Schwerte, Germany) were added to each medium. Cells were split twice a week by a confluence of ∼85%.

### 2.11 Tumor isolation

A fresh tissue sample of a 7 years old boy with a CCSK tumor was obtained for cell isolation as surplus tissue not needed for histopathological diagnosis. The tissue was mechanically minced and enzymatically digested using 5000 g/l liberase (Sigma-Aldrich, St.Louis, USA) and 240 Units of DNAse (Sigma-Aldrich, St.Louis, USA). After digestion, the cell suspension was filtered to remove debris and collected for further processing.

### 2.12 Flow cytometry

350,000 cells were incubated with fluorochrome-conjugated antibodies targeting the relevant cell surface antigens, including CD45 (Becton Dickinson GmbH, Heidelberg, Germany), GD2 (Becton Dickinson GmbH, Heidelberg, Germany) and 7-AAD (Becton Dickinson GmbH, Heidelberg, Germany). Isotype controls IgG2a and IgG1 were used. Data analysis was conducted using the FlowJo software. CD45-negative cells served as the reference population to determine the GD2 content. The comparative data between the isotype controls and the stained samples were normalized to mode.

### 2.13 R package GD2Viz development

To facilitate access to GD2 score computation and visualization, we developed GD2Viz, an R package containing an interactive web application built using R Shiny. The R package allows users to calculate GSL Reaction Activity Scores, predict GD2 scores, and explore precomputed datasets in an intuitive graphical interface.

GD2Viz is implemented in R (version 4.4.2) and follows Bioconductor guidelines to ensure integration within the Bioconductor ecosystem. Specifically, GD2Viz is designed to be interoperable with Bioconductor data structures, including SummarizedExperiment and objects, enabling the user to supply the own data as input not only in tab separated text files but also as R objects. For data privacy reasons, we separated precomputed RAS scores of several public RNA-seq datasets from the main app repository into a separate private github repository. This results in an extended online version of GD2Viz at http://shiny.imbei.uni-mainz.de:3838/GD2Viz which allows the user to explore these datasets in more depth. Without the presence of the dataset repository, the respective app tabs are hidden and the app allows the user only to analyze the own data. Additionally, the package ensures adherence to Bioconductor’s package structure, including unit testing, version control, and documentation standards, to promote long-term usability and maintenance. GD2Viz is available at https://github.com/arsenij-ust/GD2Viz, and https://arsenij-ust.github.io/GD2Viz/.

## 3. Results

### 3.1 SVM Kernel Evaluation

To assess the impact of different kernel functions on classification performance between NB and NT, we trained a variety of SVM models based on the TP adjusted RAS training data using various kernels, including 1) linear (vanilladot), 2) radial basis function (rbfdot), 3) polynomial (polydot), 4) hyperbolic tangent (tanhdot), 5) Laplace (laplacedot), 6) Bessel (besseldot), and 7) ANOVA (anovadot). Each kernel was evaluated across different hyperparameters, and models were compared based on precision, recall, F1-score, balanced accuracy (BA), and the number of support vectors (SVs). The number of support vectors serves as a key indicator of model complexity, as a lower SV count suggests better generalization and a reduced risk of overfitting.

The linear kernel (Figure 3a & f, polydot degree=1, which is equivalent to vanilladot) demonstrated high precision (0.99) but moderate recall (0.60), resulting in a F1-score of 0.75 and a BA of 0.80. It required only 174 support vectors, indicating an efficient, well-generalized model. Optimal performance for the RBF kernel was observed at σ=10 (Figure 3b - e), achieving an F1-score of 0.78 and a BA of 0.83 with 908 support vectors. The Laplace kernel (laplacedot) followed a similar trend to RBF, with the highest balanced accuracy (0.86) achieved at σ=10, but with an excessive 5,119 support vectors, suggesting overfitting (Figure 3g - k). The hyperbolic tangent (tanhdot) kernel performed poorly across all tested parameters, with models failing to generalize and yielding BA values close to 0.50 (Supplemental file 1). The Bessel kernel (besseldot) showed high recall (0.95) but poor precision (0.15) at σ=1 (SVM_33), leading to a low F1-score (0.26) (Figure 3l). Despite achieving a BA of 0.93, the model used only 191 support vectors. The ANOVA kernel (anovadot) displayed poor performance, with an F1-score of 0.12 and BA of 0.53, and a high number of support vectors (272), indicating poor generalization.

**Figure 3.**
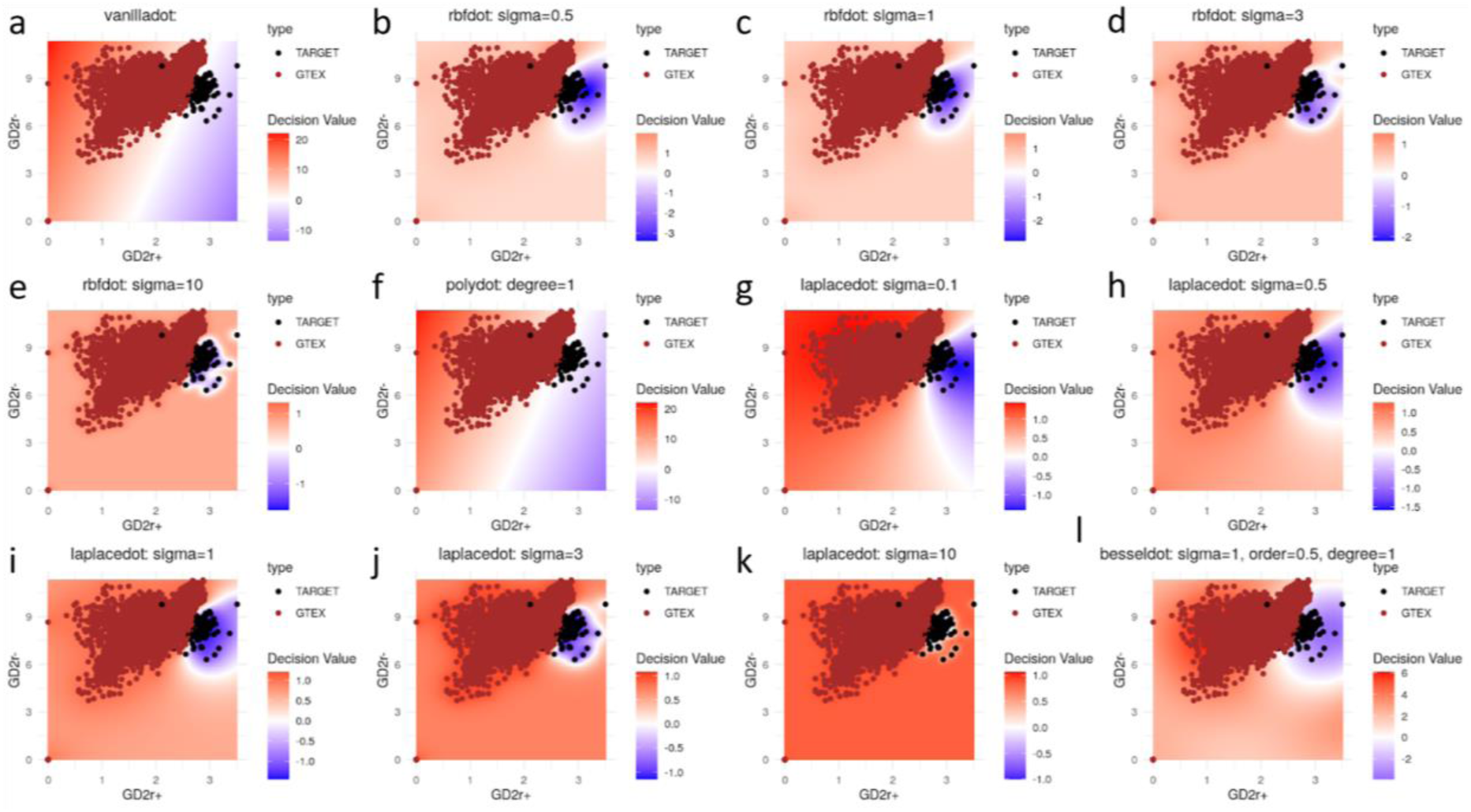
T**o**p **12 SVM models with the highest Balanced Accuracy on TP adjusted RAS training dataset.** The white area indicates the SVM hyperplane along the GD2r+ (x-axis) and GD2r-(y-axis) variables.

The results indicate that RBF and Laplace kernels with optimized sigma values (σ=10) provide the best classification performance, with the highest balanced accuracy (0.83 - 0.86). However, Laplace (σ=10) required over 5,000 SV, and RBF (σ=10) required 908 SV, which raises concerns about overfitting in regard of a two-dimensional problem with only two predictive variables. Given that the training dataset is heavily unbalanced (only 136 NB samples out of 7,548 total samples), a linear model is preferable for its generalization as it is less prone to overfitting and the interpretability in the context of the cumulative activity of GD2r+ and GD2r-reactions.

While the RBF kernel (σ=10) provides slightly higher accuracy, it does so at the cost of substantial overfitting, as evidenced by the excessive number of support vectors and by visual inspection. In contrast, the linear kernel provides a well-balanced trade-off between accuracy, generalization, and interpretability. Given the nature of the dataset and the need for explainability, the linear SVM (vanilladot) appears to be the most suitable choice for GD2 score prediction. Therefore, all GD2 score computations in the remainder of this work were performed using the linear kernel.

### 3.2 Evaluation of *ab initio* GD2r+ and GD2r-Reactions

Having showed that the linear kernel is a suitable foundation for the model, next we calculated the performance across models based on *ras*, *rasTP*, and *rasTPrec* values for the *ab initio* selected reaction set (Table 1).

**Table 1.** M**odel performance of the *ab initio* reaction set.** DR = Detection rate, DP = Detection prevalence, BA = Balanced Accuracy.

|  | *ras* | *rasTP* | *rasTPrec* |
| --- | --- | --- | --- |
| Accuracy | 0.99 | 0.99 | 0.99 |
| Kappa | 0.8 | 0.75 | 0.63 |
| Sensitivity | 0.7 | 0.6 | 0.47 |
| Specificity | 1.0 | 1.0 | 1.0 |
| Precision | 0.94 | 0.99 | 0.97 |
| Recall | 0.7 | 0.6 | 0.47 |
| F1 | 0.8 | 0.75 | 0.63 |
| Prevalence | 0.02 | 0.02 | 0.02 |
| DR | 0.0 | 0.01 | 0.008 |
| DP | 0.0 | 0.01 | 0.009 |
| BA | 0.85 | 0.8 | 0.74 |

The result suggests a strong predictive ability of the *ab initio* reaction set across *ras*, *rasTP*, and *rasTPrec*, particularly given the high balanced accuracy (0.85, 0.8, and 0.74). Accuracy and specificity are very high, while the detection rate (DR) and detection prevalence (DP) are very low, which is expected due to the imbalanced datasets. However, the decreasing trends in recall (0.7, 0.6, 0.47) and F1-score (0.80, 0.75, 0.63) indicate that the topological adjustment applied in *rasTP* and *rasTPrec* models alters the decision boundary, leading to a more conservative classification.

#### Permutation-Based Evaluation of Model Performance

To assess the robustness of the *ab initio*-defined GD2r+ (R05946, R05940) and GD2r-(R05939, R05948, R05947, R05941) reactions, a permutation test with 10,000 random permutations of GSL reaction labels was performed using a SVM with linear kernel on the *rasTP* values of the training dataset. The empirical p-values derived from these permutations were used to evaluate the significance of the model’s quality across BA, F1-score, precision and recall. The results of the *ab initio* model indicate strong classification ability, with a balanced accuracy (BA) of 0.80 (p = 0.0006), F1-score of 0.75 (p < 0.0001), precision of 0.99 (p = 0.0007), and recall of 0.60 (p = 0.0006). These p-values suggest that the *ab initio* model significantly outperforms a random reaction classifier, providing strong evidence for its predictive capability.

#### Alternative Reaction Combinations and Performance Evaluation

To further validate the *ab initio* GD2r+ and GD2r-reaction selection, 16 alternative models were constructed by modifying GD2r+ and GD2r-reaction compositions based on the position in the KEGG *glycosphingolipid biosynthesis - ganglio series* pathway. The performance of these models was evaluated using the same quality metrics (Supplemental file 1).

The results demonstrate that certain reaction combinations maintain high classification performance, whereas others fail to distinguish between NB and NT samples effectively. The highest performing alternative model (r10) composed of R05946, R05940 for GD2r+ and R05939, R05941, R05948, R05947, R05938, and R05956 for GD2r-achieved a BA of 0.81, F1-score of 0.74, precision of 0.93, and recall of 0.62 and is therefore comparable to the *ab initio* model performance (r1).

The reaction sets r2 to r9 and r11 showed a moderate classification performance with BA values ranging from 0.72 to 0.79 (median BA = 0.76), a median F1-score of 0.68, and a median recall of 0.5. These sets share the definition of reactions R05946, and R05940 as GD2r+, which seems to be the optimal combination for GD2r+.

Several reaction sets (r12 - r17) exhibited BA of 0.5, suggesting random classification performance. Based on reaction set r16, the model demonstrated a BA value of 0.57, showing marginal improvement. Notably, these results suggest that using only one reaction as GD2r+ (whether it is R05946 or R05940, which are the direct upstream reactions of GD2) or only R05948 (the first downstream reaction of GD2) as GD2r-reaction, is not enough for a good classification model.

Overall, the *ras*, *rasTP*, and *rasTPrec*-based models maintain high precision, BA, and overall reliability, reinforcing the robustness of the *ab initio* reaction set across different evaluation criteria. The results confirm that the *ab initio* GD2r+ and GD2r- reactions are robust classifiers with statistically significant predictive power. While alternative reaction combinations can maintain moderate classification performance, models lacking key reactions demonstrate a sharp decline in balanced accuracy.

### 3.3 GD2 Score Evaluation of Public Transcriptome Datasets

To investigate the distribution of GD2 scores across various transcriptome datasets, we computed the GD2 scores using the *ab initio* SVM model with linear kernel. For each dataset, the GD2 scores were derived from *ras*, *rasTP*, *rasTPrec* values. Figure 4 presents heatmaps of median GD2 scores for five RNA-seq datasets, grouped by cancer type and healthy human tissues: Figure 4a & b depict the TCGA tumor and normal tissue samples, c, d, e, and f visualize the GTEx, TARGET, St. Jude Cloud, and the CBTTC from Pediatric Brain Tumor Atlas datasets. To make the values visually comparable between the heatmaps, the color scales were standardized across the subplots. The *ras*, *rasTP*, and *rasTPrec* values are comparable across the datasets due to the size factor normalization step applied between the training data and the used datasets during prediction.

**Figure 4.**
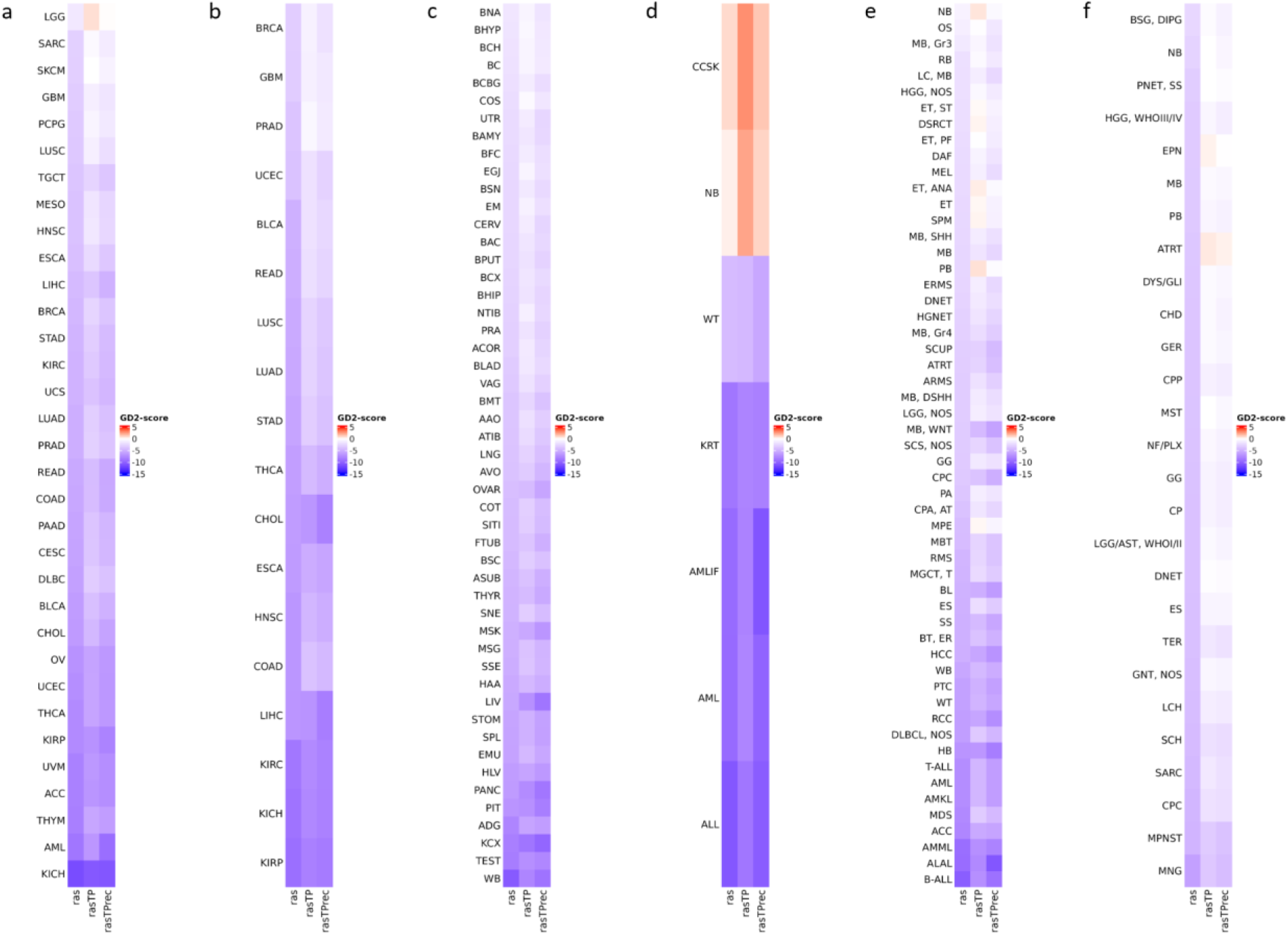
V**a**riability **of the predicted GD2 scores across cancer types for five RNA-seq datasets.** The illustrated values are the median GD2 scores per cancer type and healthy human tissues. GD2 scores were calculated on GD2r+ and GD2r- variables based on raw RAS, RAS adjusted by TP (*rasTP*), and recursive TP RAS adjustment (*rasTPrec*) values. a) TCGA tumor samples (including primary tumor, metastatic, and recurrent tumor sample types). b) TCGA normal tissue samples. c) GTEx normal tissue samples. d) TARGET tumor samples. e) St. Jude Cloud tumor samples. f) Pediatric Brain Tumor Atlas CBTTC cohort tumor samples. The *ras*, *rasTP*, and *rasTPrec* values are comparable across the subfigures. Abbreviations can be found in Supplemental file 2.

Across TARGET, St. Jude Cloud, and CBTTC datasets, NB consistently exhibited very high GD2 scores in the median. Notably, clear cell sarcoma of the kidney (CCSK) also showed a remarkably high GD2 score. Other pediatric tumors, such as Ependymoma (ET/ EPN), high-grade glioma (HGG), diffuse intrinsic pontine glioma (DIPG) and certain medulloblastoma (MB) subtypes, also showed elevated GD2 scores, suggesting GD2-expression. The median GD2 scores of certain TCGA cancer types, such as gliomas (LGG, GBM), SARC, and SKCM, were near to the hyperplane and indicate a possible variability in the distribution of GD2 scores. This variability suggests that GD2 expression may be influenced by tumor subtype, or molecular characteristics, which we further investigated in the section “GD2 Score Heterogeneity Across TCGA Cancer Subgroups”.

Both TCGA and GTEx normal tissue samples generally exhibited low GD2 scores, reinforcing the concept that GD2 expression is largely tumor-specific. Some brain-associated normal tissues from GTEx displayed higher GD2 scores than other normal tissues but were notably lower than those of NB. The GD2r- values of normal brain tissue samples were found to be higher than those of NB, consistent with the fact that in normal brain tissue, simple gangliosides are metabolized into more complex species. In contrast, NB exhibits a stronger accumulation of GD2, indicating a distinct ganglioside metabolism (Supplemental figure 1).

The three GD2 score models (*ras*, *rasTP*, and *rasTPrec*) yielded comparable distributions across all datasets, reinforcing the robustness of our scoring methodology. *rasTPrec* appeared to further refine classification by reducing noise in certain cancer types while maintaining overall consistency with the other transformations.

The GD2 score expression analysis highlights distinct GD2 expression patterns across various tumor and normal tissue types, with pediatric neuroectodermal tumors, sarcomas, and gliomas showing the highest GD2 scores. We detected a general absence of GD2 expression in normal tissues. Furthermore, the consistency between *ras*, *rasTP*, and *rasTPrec* values suggests that the GD2 scoring method is applicable across diverse transcriptomic datasets.

CCSK is predicted to express a high amount of GD2 based on our score. To validate this prediction, we analyzed GD2 expression by flow cytometry in a CCSK tumor sample and compared the results with two NB cell lines. As shown in Figure 5, the CCSK tumor exhibited GD2 expression higher than the SH-SY5Y cell line and comparable to the CHP-134 cell line, which is known for its high GD2 expression ^37^. These results validate the accuracy of our scoring method in identifying GD2-positive tumors.

**Figure 5.**
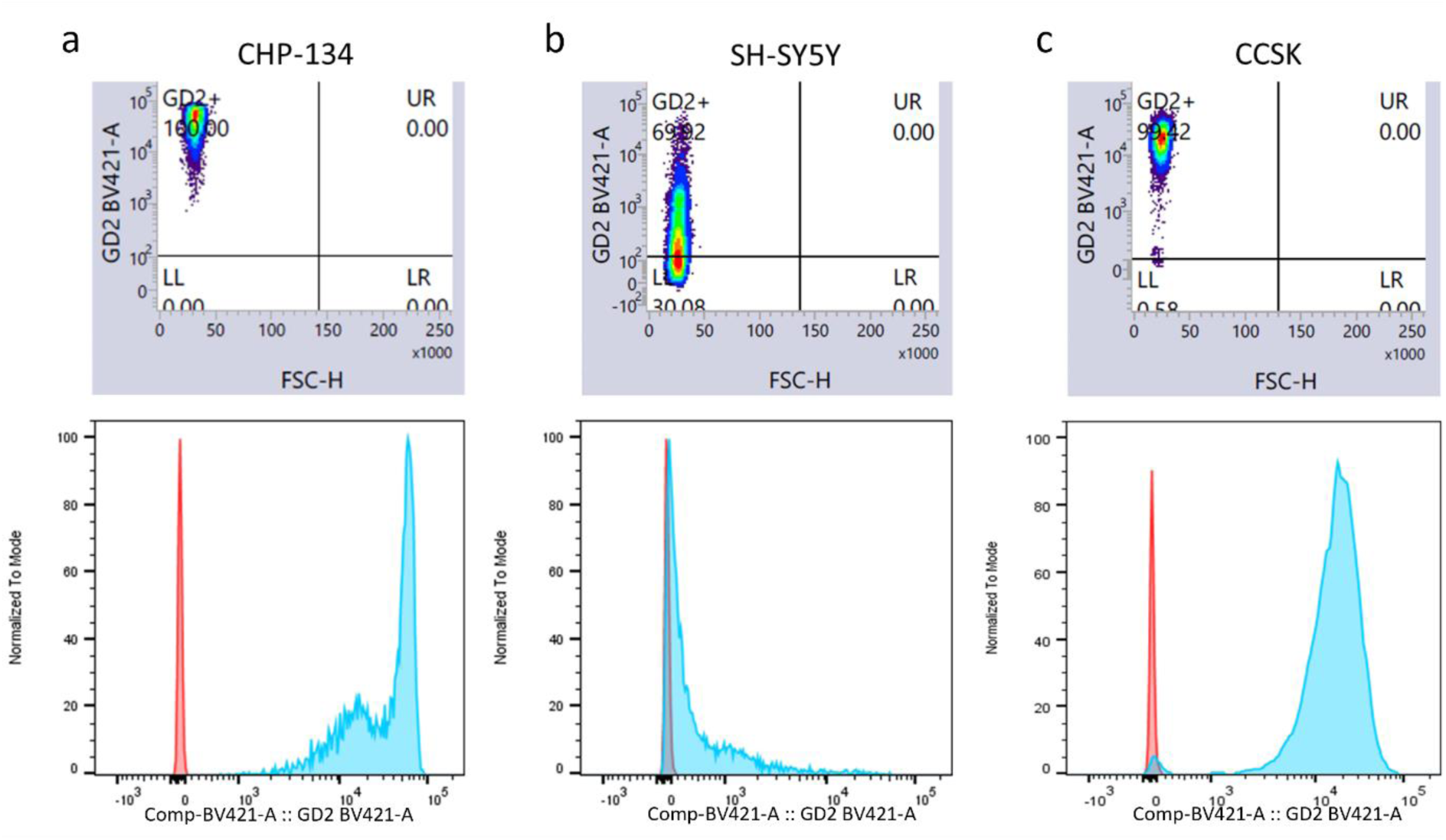
C**C**SK **expresses high levels of GD2.** GD2 expression was analyzed by flow cytometry in the two NB cell lines CHP-134 (a) and SH-SY5Y (b) and in one CCSK tumor sample (c). In the lower panel, isotype control is showed in red and GD2 expression in blue.

### 3.4 Comparison of the GD2 Score with the Two-Gene Signature

To further validate the GD2 score, we compared it to an established two-gene signature from *Sorokin et al.* 2022 across six independent RNA-seq datasets, assessing its performance in distinguishing GD2 expression levels across cancer types in accord with the literature’s expected outcome (Figure 6 and Supplemental file 3). The datasets include MB subtypes from GSE203174 and CBTTC (Figure 6a, b), H3 wildtype and mutant HGG, and diffuse midline gliomas (DMG) (Figure 5c, d), GN and NB from GSE147635 (Figure 6e), and GD2-high/low parental Kelly cells (NB) stained with GD2-APC antibody (Figure 6f).

**Figure 6.**
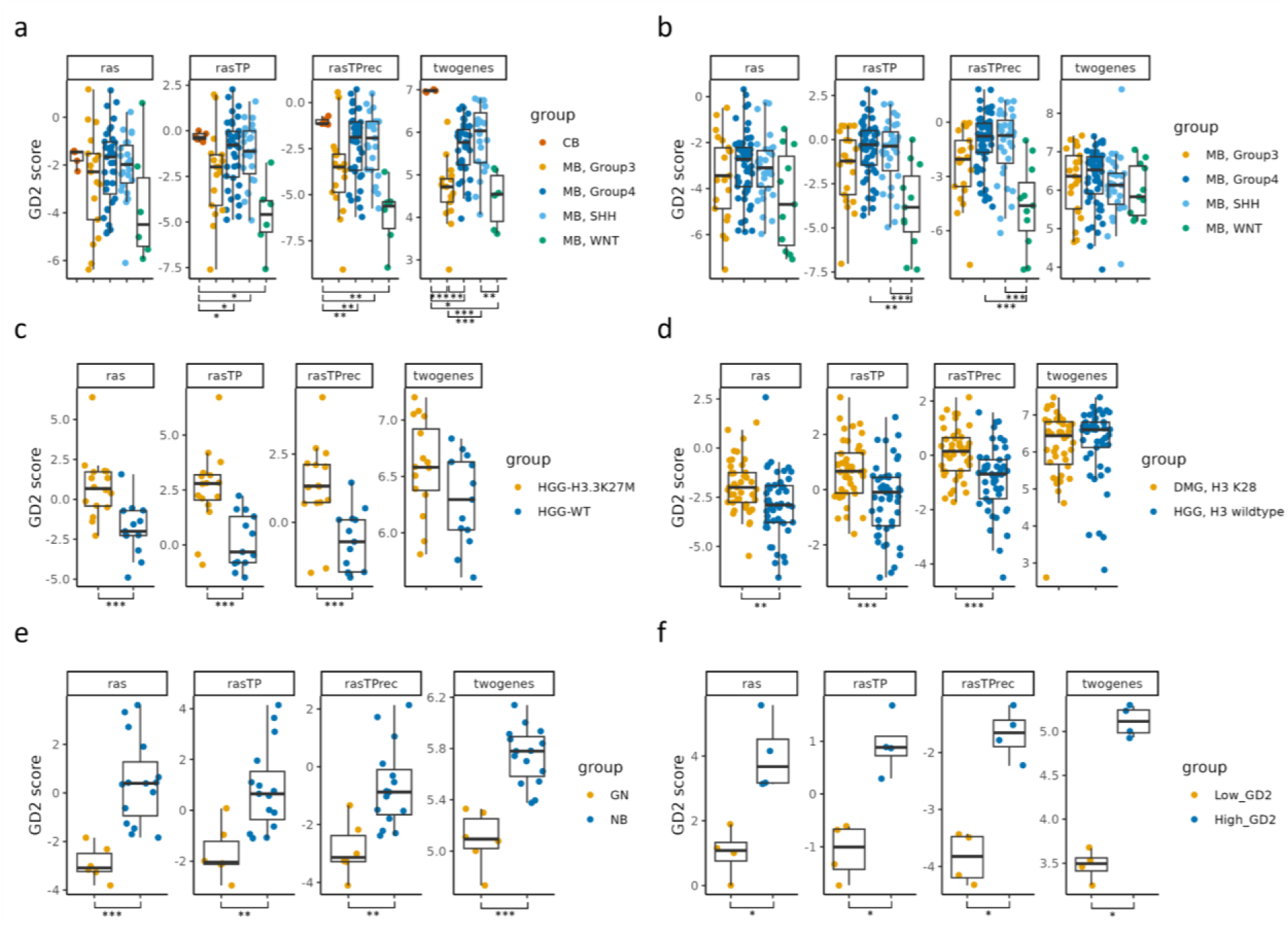
V**a**lidation **of the predicted GD2 scores for six RNA-seq dataset and comparison with the two-gene signature.** Boxplots visualizing the GD2 scores calculated on GD2r+ and GD2r- variables based on raw RAS, RAS adjusted by TP (*rasTP*), and recursive TP RAS adjustment (*rasTPrec*) values, as well as the two-gene signature, for six RNA-seq datasets, namely: a) medulloblastoma (MB) subtypes and normal cerebellum samples (CB) from GSE203174. b) medulloblastoma (MB) subtypes from the CBTTC cohort. c) high-grade glioma (HGG) samples with an H3K27M alteration and H3-3 wildtype (WT) from GSE117446. d) diffuse midline glioma with an H3K27M alteration (H3K28 is a synonym for H3K27M) and high-grade glioma with H3-3A wildtype from the CBTTC cohort. e) ganglioneuroma (GN) and neuroblastoma (NB) from GSE147635. f) parental Kelly cells (neuroblastoma) stained with GD2-APC antibody and sorted in GD2-low and GD2-high populations from GSE180514. *** = adj. p ≤ 0.001; ** = 0.001 < adj. p ≤ 0.01; * = 0.01 < adj. p ≤ 0.05; not significant = adj. p > 0.05 (not illustrated). The sample numbers per group, p-values, and effect sizes are documented in Supplemental file 3.

Across all datasets, the *ras*, *rasTP*, and *rasTPrec* derived GD2 scores and the two-gene signature demonstrated comparable patterns. Expected differences across tumor subtypes and experimental groups were identified by both methods. Notably, while the two-gene signature indicated the highest GD2 expression in normal cerebellum (CB), the GD2 scores indicated a lower GD2 expression compared to some high-GD2 score samples in MB group 3, group 4, and SHH (Figure 6a). Also, the GD2 score could better discriminate the MB subgroups of the CBTTC cohort and between H3 wildtype and mutant samples of the GSE117446 and CBTTC datasets in comparison to the two-gene score (Figure 6b, c, and d).

In both GSE203174 (Figure 6a) and CBTTC cohort (Figure 6b), MB subtypes exhibited distinct GD2 score distributions. The WNT subtype consistently displayed lower scores, while most of the samples of group 3, group 4, and SHH MB had significantly higher GD2 scores. This is in agreement with prior knowledge of GD2 expression patterns, where both SHH and group 4 express GD2, whereas group 3 is more diverse ^12^.

In GSE117446 (Figure 6c) and CBTTC cohort (Figure 6d), samples harboring the H3.3K27M alteration had significantly higher GD2 scores than their H3 wildtype counterparts, supporting the hypothesis that GD2 expression is associated with this specific alteration ^38,39^.

In GSE147635 (Figure 6e), neuroblastomas exhibited significantly higher GD2 scores than ganglioneuromas, aligning with the expected GD2 expression profile of these tumors ^12,40^. In GSE180514 (Figure 6f), the GD2 score and the two-gene signature confirmed a significant difference in GD2 scores between GD2-high and GD2-low Kelly cells.

Overall, these findings highlight that the GD2 scores based on *ras*, *rasTP*, *rasTPrec* can capture GD2-associated expression differences across multiple datasets and tumor types, similar to the performance of the previously described two-gene signature. However, the GD2 score differs in several key factors, namely the lower GD2 expression in brain tissues relative to some MB samples, and the different patterns of GD2 expression across subtypes.

### 3.5 GD2 Score Heterogeneity Across TCGA Cancer Subgroups

#### mRNA-Based Subgroups

To investigate the variability in GD2 expression among different molecular subtypes of cancer, we exemplarily identified five TCGA cancer types that showed differences in GD2 scores across their subtypes using published mRNA-based classification for subtype annotation (Figure 7). These findings build upon the observations from Section 3.3, where cancers such as gliomas (LGG, GBM), SARC, and SKCM exhibited median GD2 scores near the hyperplane, suggesting intratumoral heterogeneity in GD2 expression.

**Figure 7.**
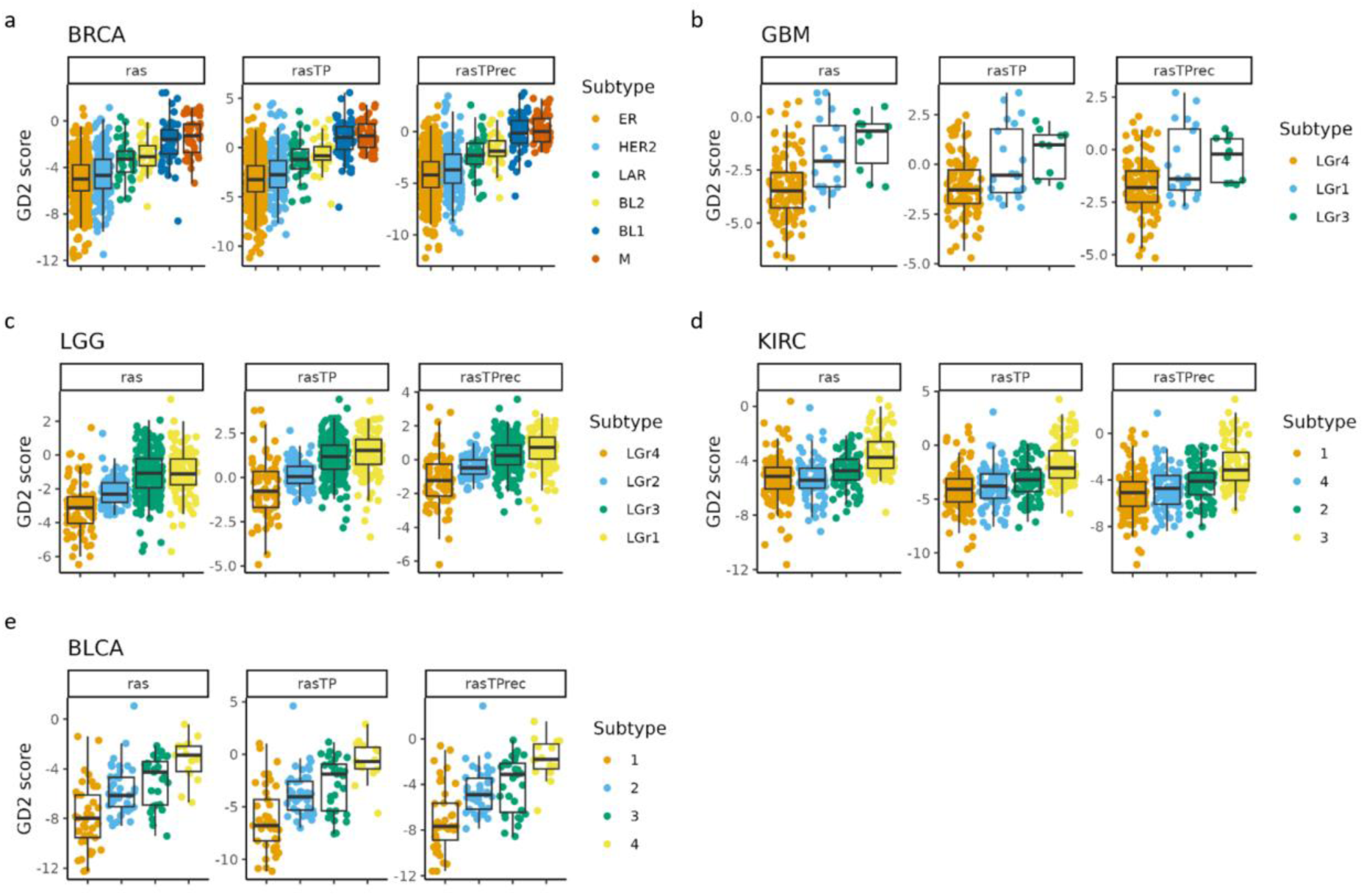
G**D**2 **score variability across mRNA-based cancer subtypes indicates subgroup specific GD2 expression patterns.** Boxplots visualizing the GD2 scores calculated on GD2r+ and GD2r- variables based on raw RAS, RAS adjusted by TP (*rasTP*), and recursive TP RAS adjustment (*rasTPrec*) values of five TCGA projects: a) TCGA breast cancer (BRCA). The subgroup labels were assigned according to the reported results of *Lehman et al.* 2016 ^29^. BRCA subtypes: ER = estrogen receptor; HER2 = human growth factor receptor 2; LAR = luminar androgen receptor; BL1 and BL2 = basal-like subtypes; M = mesenchymal. b) TCGA glioblastoma multiforme (GBM). c) TCGA low-grade glioma (LGG). GBM and LGG subtypes were assigned based on reported gene expression clusters in *Ceccarelli et al.* 2016 ^41^. d) TCGA kidney clear cell carcinoma (KIRC). KIRK RNA expression subtypes based on reported clusters in ^42^. e) TCGA bladder urothelial carcinoma (BLCA). BLCA subtypes were assigned according to ^43^. The sample numbers per group, and Dunn’s p-values, are documented in Supplemental file 3.

Among BRCA subtypes (Figure 7a), the mesenchymal (M) and basal-like (BL1) subgroups exhibited the highest GD2 scores, followed by BL2 and LAR-positive subtypes. In contrast, estrogen receptor-positive (ER) and HER2-enriched subtypes showed in median lower GD2 scores but a higher variance across samples, suggesting that GD2 expression is enriched in basal-like and mesenchymal BRCA subtypes rather than luminal or hormone receptor-driven tumors. GBM and LGG exhibited distinct subtype-dependent differences in GD2 scores. In GBM (Figure 7b), LGr3 had the highest GD2 scores, while LGr4 had moderate to low scores. Notably, LGr1 has a mixed expression pattern, more visible in *rasTP* and *rasTPrec* with some samples having exceptionally high GD2 scores. The GD2 score pattern across the mRNA subgroups of GBM is consistent with LGG (Figure 7c). Here, LGr1 and LGr3 exhibited higher GD2 scores compared to LGr2 and LGr4. GD2 scores in KIRC subtypes (Figure 7d) were comparable across the mRNA subgroups 1, 2, and 4, indicating a GD2 score below zero in all three models. Only group 3 showed a remarkably higher score, suggesting the existence of a separated group of samples that may express a higher amount of GD2. BLCA exhibited heterogeneity in GD2 scores across its subtypes (Figure 7e). Subgroup 4 had the highest GD2 scores, followed by subgroups 2 and 3, whereas subgroup 1 had lower scores but a broader distribution of samples.

These findings demonstrate that the predicted GD2 expression varies across several cancer subtypes based on mRNA characterization, particularly observed in BRCA, GBM, LGG, KIRK, and BLCA. The drivers of these variations have to be further investigated and integrated with other data types like mutations, fusions, and copy number alterations. Nevertheless, these results underscore the importance of tumor-specific and subtype-specific GD2 evaluation, which may have implications for GD2-targeted immunotherapy strategies.

#### GD2-Associated Copy Number Alterations

By integrating gene-level copy number alterations (CNAs) with the GD2 score, we observed possible associations between high-level amplification of cancer-related genes in three TCGA projects: GBM, SARC, and LUAD (Figure 8). Using processed CNA data, we assessed whether differences in CNA status were associated with significant changes in the GD2 score. For each gene within defined subgroups, we employed the Kruskal-Wallis test to compare GD2 scores across the CNA status levels (homozygous deletion, hemizygous deletion, no change, gain, and high-level amplification). Genes with an adjusted p-value below 0.05 and an η^2^ exceeding 0.06 were considered to have a statistically significant association. The left column of Figure 8 displays boxplots of GD2 scores across histological or molecular subtypes within each TCGA project. The coloring represents the CNA status of *B4GALNT1*, the gene coding the enzyme metabolizing GD3 to GD2. The right column presents GD2 scores stratified by the CNA status for several genes of interest in one of the cancer subtypes per project.

**Figure 8.**
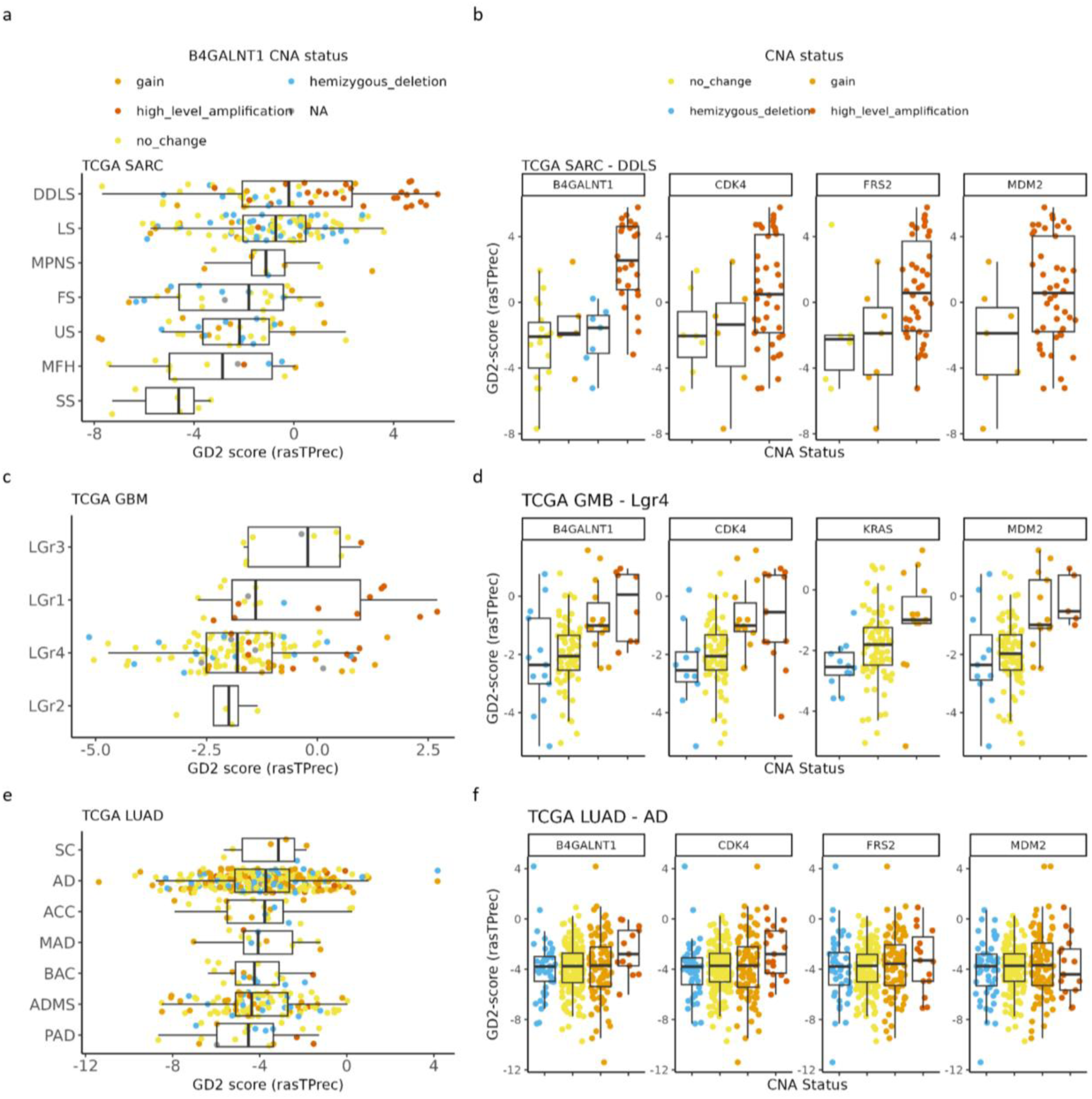
B***4***GALNT1 **amplification is associated with high GD2 scores in dedifferentiated liposarcoma.** GD2 score based on recursive TP RAS adjustment (*rasTPrec*) values of three TCGA projects grouped by subtypes (left column) and selected genes grouped by copy number alteration (CNA) status for three subgroups (right column). a) TCGA sarcoma (SARC), DDLS = Dedifferentiated liposarcoma, LS = Leiomyosarcoma, MPNS = Malignant peripheral nerve sheath tumor, FS = Fibromyxosarcoma, US = Undifferentiated sarcoma, MFH = Malignant fibrous histiocytoma, SS = Synovial sarcoma. b) CNA of *B4GALNT1*, *CDK4*, *FRS2*, and *MDM2* from TCGA dedifferentiated liposarcoma (DDLS). c) TCGA glioblastoma multiforme (GBM). d) CNA of *B4GALNT1*, *CDK4*, *KRAS*, and *MDM2* from TCGA GBM LGr4. e) TCGA lung adenocarcinoma (LUAD), SC = Solid carcinoma, AD = Adenocarcinoma, ACC = Acinar cell carcinoma, MAD = Mucinous adenocarcinoma, BAC = Bronchiolo-alveolar carcinoma, ADMS = Adenocarcinoma with mixed subtypes, PAD = Papillary adenocarcinoma. f) CNA of *B4GALNT1*, *CDK4*, *FRS2*, and *MDM2* from TCGA lung adenocarcinoma.

The GD2 score varied significantly across TCGA sarcoma subtypes with the highest GD2 scores found in dedifferentiated liposarcoma (DDLS) (Figure 8a). For each subtype, we identified genes, that were significantly associated with the GD2 score as described above. In DDLS, the analysis revealed that CNA levels in *B4GALNT1* (adj. p-value = 0.02, and large effect size = 0.5) was significantly associated with differences in the GD2 score (Figure 8b). Specifically, samples exhibiting high-level amplifications of *B4GALNT1* tended to have higher GD2 scores compared to those with copy number gain, no change or deletion. These results are logical, because *B4GALNT1* encodes one of the key enzymes in GD2 biosynthesis. In contrast, the adjusted p-values of *CDK4* (adj. p-value = 0.6, and moderate effect size = 0.07), *FRS2* (adj. p-value = 0.6, and moderate effect size = 0.09), and *MDM2* (adj. p-value = 0.6, and small effect size = 0.06) were not significant.

Among GBM mRNA subgroups, significant heterogeneity in GD2 scores was already discussed in section 3.5. In GBM LGr4, 804 copy-number altered genes were identified to have a significant association. Focusing on CNA status of *B4GALNT1*, we detected significant associations with the GD2 score in LGr4 (adj. p-value = 0.02, and large effect size = 0.16), as well as in *CDK4* (adj. p-value = 0.02, and large effect size = 0.15), and *MDM2* (adj. p-value = 0.02, and large effect size = 0.16) (Figure 8d). Notably, among significant genes, *KRAS* (adj. p-value = 0.048, and moderate effect size = 0.09) and ST8SIA1 (adj. p-value = 0.03, and moderate effect size = 0.1), another relevant gene in the GD2 biosynthesis, showed an association between CNV gain status and GD2 score. While LGr1 samples with the highest GD2 score also have high-level *B4GALNT1* amplifications, statistically this was not significant with an adjusted p-value of 0.5 and a large effect size of 0.27, due to low sample sizes in this group. These results indicate that, similar to DDLS, *MDM2* or *CDK4* could serve as a surrogate biomarker for samples with a *B4GALNT1* amplification in GBM, which leads to a high GD2 score.

LUAD cases, stratified by histological subtypes, also exhibited distinct GD2 score profiles. In LUAD subtype adenocarcinoma (AD) (Figure 8f), samples with high-level amplification in *B4GALNT1*, *CDK4*, *FRS2*, and *MDM2* were observed but were not significantly associated with GD2 score differences (adj. p-values: 0.49, 0.48, 0.86, 0.97 and effect sizes: 0.007, 0.01, -0.004, -0.007). In contrast to SARC and GBM, samples with amplifications or gains in *B4GALNT1*, *CDK4*, *FRS2*, and *MDM2* did not show elevated GD2 scores, suggesting that other, currently unknown, key drivers of GD2-associated phenotypes in lung adenocarcinoma might exist but might be missed by standard transcriptomic assays.

In summary, our analysis reveals that the GD2 score is significantly modulated by CNAs in *B4GALNT1* which could be due to a co-amplification with key oncogenic genes across multiple cancer types. The significant associations of *B4GALNT1* and the overlap with established oncogenic drivers localized on the same chromosome from OncoKB, notably *CDK4* and *MDM2*, underscore the potential mechanistic link between these CNAs and GD2 expression.

#### Higher GD2 Score in Squamous Cell Carcinoma

We observed differences in GD2 score across adenocarcinoma and squamous cell carcinoma (SCC) in the following TCGA projects: esophageal carcinoma (ESCA), head and neck squamous cell carcinoma (HNSC), cervical and endocervical cancer (CESC), and the combined cohort of lung squamous cell carcinoma (LUSC) and LUAD (Figure 9). In TCGA-ESCA, GD2 scores were significantly higher in SCC compared to AD across all three models (*ras*-model p-value = 1.5e-13, *rasTP*-model p-value = 1e-16, *rasTPrec*-model p-value = 8.6e- 18), suggesting an association between GD2 expression and squamous histology in esophageal tumors (Figure 9a). The GD2 scores were relative similar across the SCC subtypes of HNSC, while basaloid SCC exhibited slightly lower GD2 scores (Figure 9b). The GD2 score pattern of HNSC SCC was similar to SCC of ESCA tumors. GD2 scores were consistently elevated in SCC, with keratinizing SCC and large cell nonkeratinizing SCC showing higher scores compared to adenocarcinoma subtypes (Figure 9c). In LUSC, GD2 scores of SCC, keratinizing SCC, and basaloid SCC were higher than LUAD subgroups, including AD, acinar cell carcinoma, papillary AD, mucinous AD, bronchiolo-alveolar carcinoma, and AD with mixed subtypes, further emphasizing a potential correlation between GD2 expression and squamous differentiation in lung tumors (Figure 9d). Across all four cancer types analyzed, GD2 scores were consistently higher in SCC compared to AD. Further analysis is required to identify possible molecular, clinical or histological drivers for the observed higher GD2 scores in squamous cell carcinoma.

**Figure 9.**
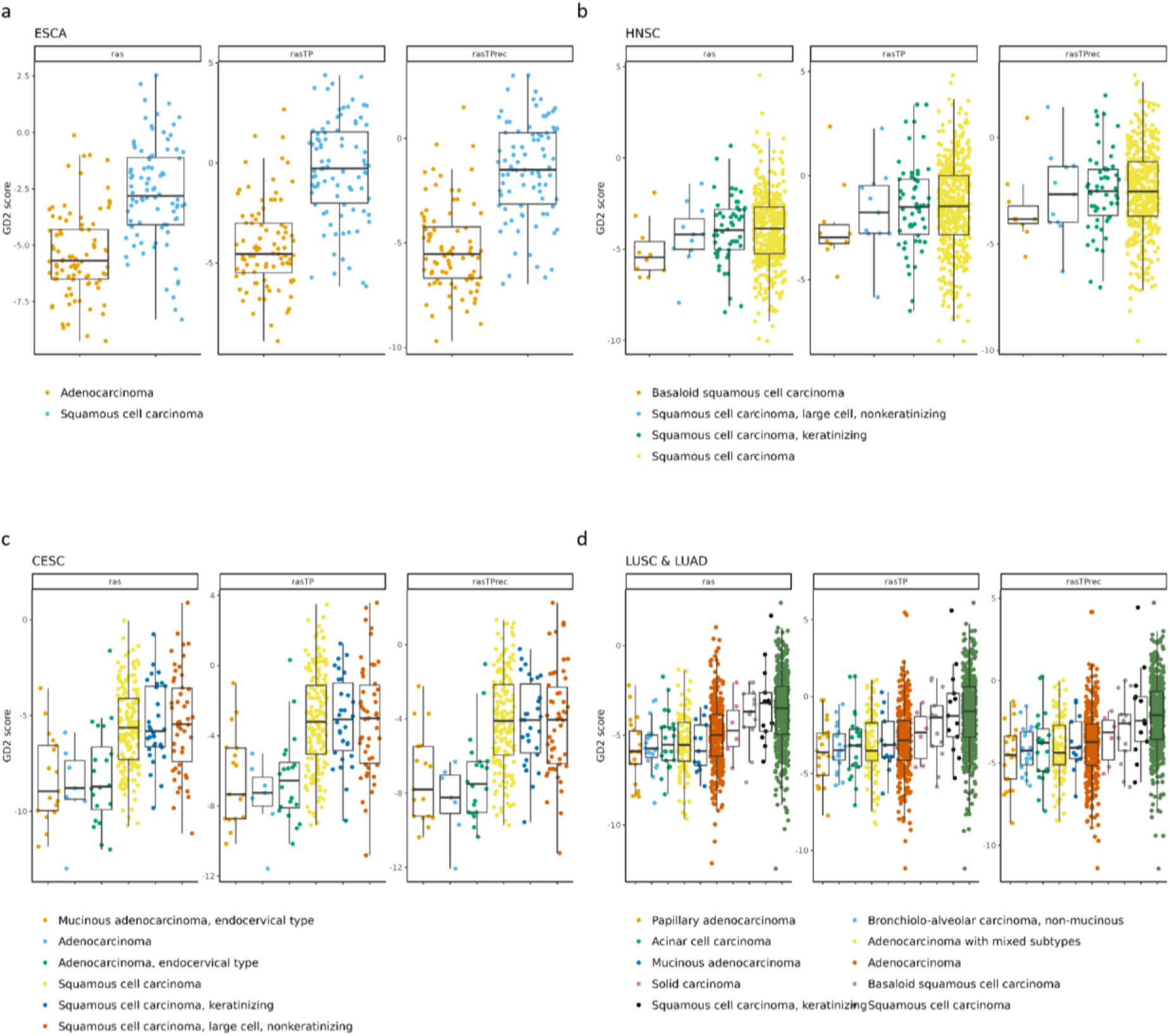
S**q**uamous **cell carcinoma exhibit higher predicted GD2 scores compared to adenocarcinoma.** Boxplots visualizing the GD2 scores calculated on GD2r+ and GD2r- variables based on raw RAS, RAS adjusted by TP (*rasTP*), and recursive TP RAS adjustment (*rasTPrec*) values of four TCGA projects: a) TCGA esophageal carcinoma (ESCA), AD (n=87), squamous cell carcinoma (n=86). b) TCGA head & neck squamous cell carcinoma, basaloid SCC (N=10), SCC large cell nonkeratinizing (n=11), SCC keratinizing (n=54), SCC (n=442). c) TCGA cervical & endocervical cancer (CESC), mucinous AD (n=17), AD (n=7), AD endocervical (n=20), SCC (n=170), SCC keratinizing (n=30), SCC large cell nonkeratinizing (n=50). d) TCGA lung squamous cell carcinoma (LUSC) and TCGA lung adenocarcinoma (LUAD), papillary AD (n=22), bronchiolo-alveolar carcinoma (n=19), Acinar cell carcinoma (n=22), AD with mixed subtypes (n=106), mucinous AD (n=13), AD (n=311), solid carcinoma (n=6), basaloid SCC (n=13), SCC keratinizing (n=13), SCC (n=464).

### 3.6 Using the GD2 Score Methodology via the R Shiny GD2Viz Package

In order to facilitate access to the aforementioned methodology for scientists and physicians, we developed the GD2Viz R/Shiny package. This offers the possibility to carry out the necessary process steps, starting with the calculation of the RAS values for the GSL pathway from a non-normalized count matrix, up to the GD2 score prediction in one’s own R environment. The primary feature of GD2Viz is an interactive application that enables any researcher to effortlessly evaluate their data sets in terms of the GD2 score.

The app consists of three tabs (Figure 10): The “Public Datasets” tab visualizes the precomputed GD2 score across six major RNA-seq datasets: “TCGA Tumor samples”, “TCGA normal samples”, “GTEx”, “TARGET”, “St. Jude Cloud”, and the “CBTTC” dataset from the Pediatric Brain Tumor Atlas. The user can choose between an interactive scatterplot, boxplot, or violin plot view, as well as explore the slightly different results by changing the SVM model settings, e.g. use raw, ranged, or scaled RAS values as data for model training or choose the preferred RAS adjustment method. Additional features, like grouping and highlighting by an experimental variable, allow further easy exploration of the datasets (Figure 10a).

**Figure 10.**
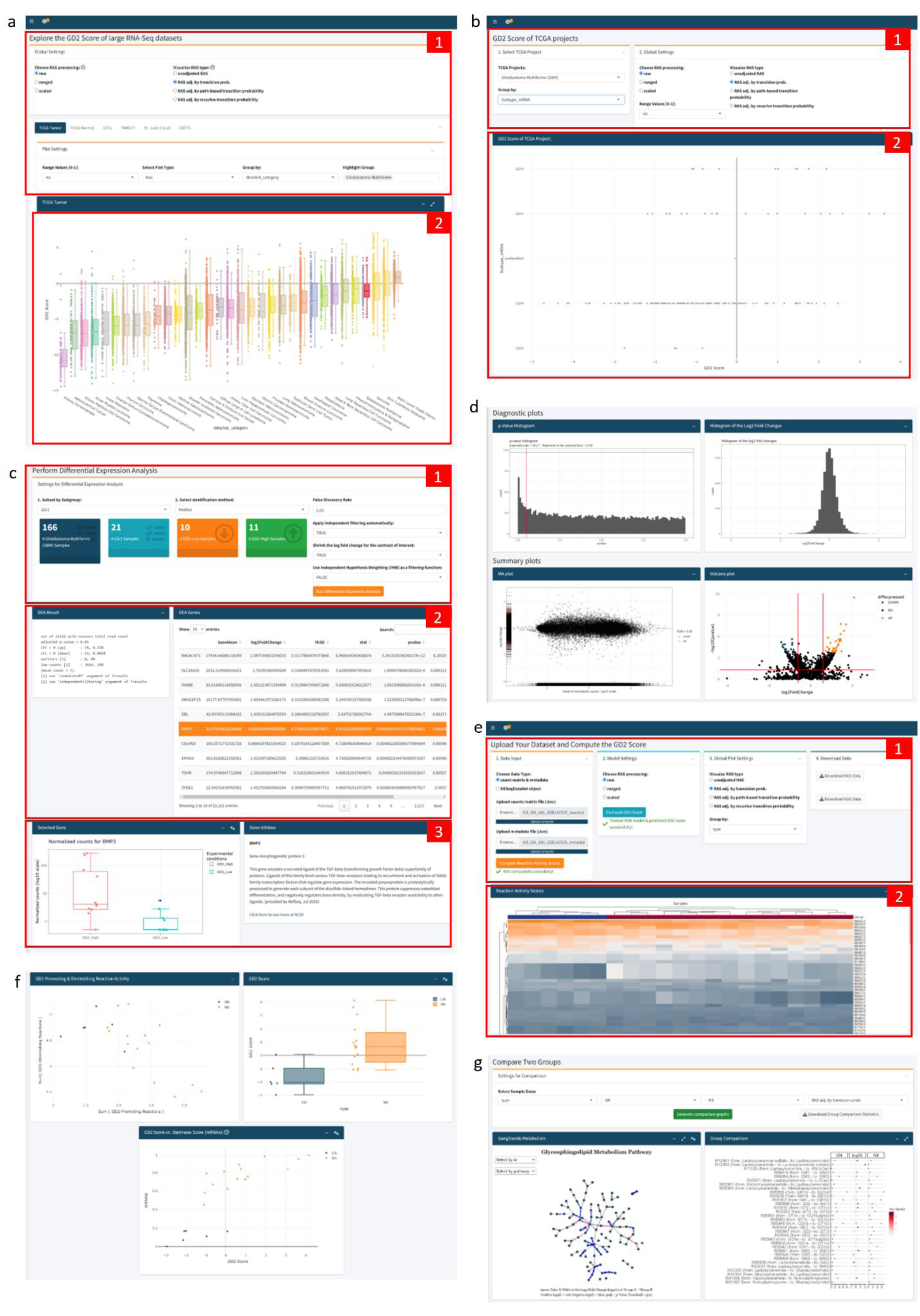
A**n**notated **Screenshots of GD2Viz Shiny app.** a) First Tab contains the model and plot settings (1) and the GD2 score visualization of the selected dataset (2). b) The top part of the second tab contains the selections for the TCGA project, the experimental variable, and the model settings (1). (2) shows the GD2 score of the TCGA project grouped by the selected molecular subtype. c) Depicts the DEA part of the second tab, made up of the settings for the DEA (1), the summary of the DEA and statistics for each gene (2), and the gene expression and information of the selected gene (3). d) Illustration of the diagnostic and summary plots of the DEA. e) The top section of the third tab contains the file upload functionality, model and plot settings, as well as the possibility to download the computed RAS and GD2 score values (1). (2) shows the heatmap of the adjusted RAS. f) Plots of GD2r+ and GD2r- variables, the GD2 score grouped by an experimental variable of the uploaded dataset, and the GD2 score against the stemness score. g) Group comparison section of the third tab, visualizing the log2 fold-change of all GSL network reactions. Screenshots were taken of the GD2Viz app version 0.1.1.

The second tab focuses on single projects within the TCGA project (Figure 10b). It allows to analyze the individual tumor datasets in terms of clinical variables or molecular subtypes. Similar to the first tab, the model parameters can be selected. Besides the GD2 score visualization, the tab provides the possibility to run a differential expression analysis (DEA) for the entire project or a specific subset of samples (Figure 10c). For the DEA, the GD2 score is used for stratification of the samples in two (GD2-high vs. GD2-low) or three (GD2-high, GD2- medium, GD2-low) groups. The results display the overall result of the DEA, various diagnostic and summary plots, such as MA- and interactive volcano plots, a searchable table of significant genes, and additional gene information of the selected gene (Figure 10c - d).

The third tab enables users to upload their datasets and compute the RAS values and the GD2 score (Figure 10e). The input can be a raw count matrix, which is a commonly obtained object during RNA-seq analysis, and a metadata file in .tsv format from an RNA-seq experiment or a DESeqDataSet object in .rds format. Users can compute adjusted and raw RAS values, change model settings, and predict the GD2 scores. Interactive visualization tools include adjustable heatmaps of RAS values, scatter plots showing GD2-promoting vs. GD2- diminishing reactions, and plots of the predicted GD2 scores. Additional plots illustrate the GD2 score against selected genes or stemness scores (Figure 10f). A comparison section provides a log2 fold-change analysis of Glycosphingolipid metabolism and a detailed ganglioside pathway analysis including a network visualization (Figure 10g). The final section supports differential gene expression analysis based on GD2 score stratification, similar to the functionality of the second tab.

## 4. Discussion

To predict GD2 expression based on transcriptomic data, we developed a computational pipeline integrating metabolic network modeling and machine learning. RAS were computed for the enzymatic reactions of the GSL metabolic network and adjusted by transition probabilities incorporating the network topology. Using the cumulative RAS of *ab initio* selected GD2 mitigating and promoting reactions, an SVM was trained to distinguish neuroblastoma from normal tissue samples. The SVM model provides predicted decision values rather than binary classification outputs or probabilities. Only in the case of a linear kernel and a two-class classification problem, the decision values can be interpreted as the distance of each sample from the hyperplane. These values are particularly meaningful in the context of the GD2 phenotype, which exists on a continuum between low and high expression levels. The decision values function as a quantitative proxy for the GD2 score, offering a more refined perspective than a binary classification. Therefore, the score represents the degree of association with either the GD2-positive or GD2-negative phenotype, with values closer to the hyperplane boundary indicating intermediate states. Furthermore, the two variables GD2r+ and GD2r- are related to this continuum and permit an interpretation of whether the overall activity of the ganglioside pathway is high or low, and whether the GD2 mitigating reactions dominate the GD2 promoting reactions, potentially indicating the further metabolization of GD2 rather than its accumulation.

Since the decision values of the SVM are used as a continuous GD2 score, the lower rate of identifying true positives (lower recall) is mitigated. The flexibility of thresholding ensures that sensitivity can be optimized, when necessary, by adjusting the decision boundary to capture more true positives. This adaptability allows for fine-tuning based on specific clinical or experimental requirements, ensuring a balance between recall and precision. Given that all three models exhibit strong classification performance, the selection of a single best model remains challenging, especially in the absence of direct validation of GD2 expression in the training dataset. The consistently high precision and balanced accuracy across models underscore the robustness of the *ab initio* reaction set. Further experimental validation using laboratory-based methods is essential to confirm our findings and enhance the clinical applicability of this approach. We were indeed able to confirm that CCSK expresses high level of GD2, how predicted by the GD2 score, by measuring GD2 on a tumor sample by flow cytometry in comparison to NB. Notably, this is the first time that GD2 expression in CCSK has been demonstrated. CCSK are the second most prevalent pediatric kidney tumor, after Wilms’ tumor and have a less favorable prognosis. New targeted approaches are urgently required to improve outcomes for patients with advanced-stage disease ^44^. Previous study suggests that CCSKs originate from a renal mesenchymal cell exhibiting a diverse array of neural markers. As a result, these cells appear prone to alterations also observed in various other neuroectodermal and neuronal tumors, which could explain the high level of GD2 expression ^45^.

When comparing GD2 scores derived from *ras*, *rasTP*, and *rasTPrec* models, we observe that *ras* and *rasTPrec* are more restrictive, leading to a higher number of samples falling below zero, whereas *rasTP* is more liberal, resulting in a greater number of samples above zero.

The GD2 score has been shown to have certain advantages over the two-gene signature, including consistency across datasets. The results suggest reliability and robustness, although the used test datasets were not batch-corrected with the training dataset, apart from the size factor normalization. Moreover, the presence of a hyperplane indicating an intermediate GD2 expression helps in the interpretation of the predicted GD2 scores. In contrast, a two-gene signature lacks uniform scaling, requiring reference samples with known GD2 expression levels to contextualize results.

One limitation of our predictive model is the multi-layered regulation of GD2 expression, which is modulated by a number of factors, including the activity of key enzymes involved in ganglioside metabolism, namely the one encoded by *ST3GAL5*, *ST8SIA1*, *B4GALNT1*, and *B3GALT4*. The activity of these enzymes can be modulated by transcriptional and posttranslational mechanisms, including phosphorylation and N-glycosylation ^46^, and is further dependent on the developmental stage ^47^. These factors contribute to the variability of GD2 expression in normal and cancerous tissues, thus adding another layer of complexity to GD2 regulation. We acknowledge that the regulation of ganglioside metabolism is a complex process involving multiple factors, which modulate the expression patterns and are not fully captured by our model.

Using the RAS-based GD2 score, we were able to reproduce known data on GD2 expression across and within tumor entities. This includes heterogeneous expression of GD2 within different pediatric tumors such as NB, MB and DMG. GD2 expression in NB varies among patient samples, with some exhibiting intermediate to low levels ^12^. This variability may be related to a more differentiated cell state ^12^. Our results align with these clinical observations, as some NB samples in both the training and test datasets exhibited intermediate to low GD2 scores. Heterogeneous and subtype-dependent GD2 expression in MB has been previously described ^12^. In accord with our results, the lowest expression of GD2 was detected in the WNT subtype and the highest variability in group 3, where some samples were GD2 negative due to the accumulation of GM3 ^12^. Further, our score indicates that in diffuse midline glioma, the H3K27M mutation is predictive of GD2 expression, as shown in previous work assessing GD2 expression by flow cytometry in pediatric high-grade gliomas with mutated or wild-type H3 ^38^.

Notably, our approach suggests low expression of GD2 in brain tissues, especially in the cerebellum, while based on the two-gene signature high expression would be expected. A low amount of GD2 in the brain and peripheral nerves is in accord with clinical data showing no on-target off-tumor toxicity in patients treated with CAR-T cells against GD2 ^48,49^. This highlights the importance of incorporating GD2-diminishing reactions into the GD2 score. GD2 serves as an intermediate glycolipid in the synthesis of complex gangliosides characteristic of the mature brain, whereas simpler gangliosides like GD2 are more typical of the developing embryonic brain ^47^.

Based on the GD2 score, subtypes of other tumor entities can also express high levels of GD2. These subtypes may reflect biologically and clinically relevant subsets. In diffuse glioma for example we observed heterogeneity of GD2 expression in previously described mRNA- based subgroups ^41^. A particularly high GD2 score was found in mRNA-based subgroups LGr1 and LGr3, both characterized by the presence of IDH mutations. The LGr4 subgroup, which includes particularly GBM samples, is highly heterogeneous, being composed of classical, mesenchymal, neural, and proneural subtypes, which may explain the high variability in GD2 score values of this cluster ^41^. In GBM, high GD2 expression has been indeed observed by using lipid analysis on tissue samples ^50^ but not all tumor samples expressed GD2. This underlines the necessity to assess GD2 expression before treating patients with anti-GD2 therapies or to identify biomarkers predicting GD2 expression. An anti-GD2 antibody-drug conjugate is currently in clinical testing for GD2 positive tumors, including glioblastoma (NCT06641908).

In breast cancer, a GD2-positive population (up to 35%) has been identified by flow cytometry analysis of human breast cancer cell lines and patient samples. This GD2-positive population has a cancer stem cells phenotype and possesses the ability to form mammospheres and initiate tumors ^51^. Our data indicates that particularly two subtypes of TNBC, the mesenchymal and the basal-like 1 subtype express GD2. Expression of GD2 in TNBC has been previously described by immunohistochemistry of frozen samples and associated with worse overall survival of patients ^52^. TNBC is very heterogeneous and it is associated with high recurrence rates, distant metastasis, and unfavorable clinical outcomes. TNBC cells lack targetable receptors; hence, targetable markers are urgently needed. GD2 expression could be used to stratify these TNBC subtypes and as targets for therapeutic interventions.

The lowest GD2 scores were observed in mRNA subgroups 1 and 2 of bladder urothelial carcinoma. These subgroups exhibit high HER2 protein levels, which are similar to the HER2 and LAR subtypes of BRCA ^43^. These also exhibited the lowest GD2 scores in median. Furthermore, markers of urothelial differentiation are also highly expressed in subgroups 1 and 2. Group 3 and 4 of BLCA correlate with the TCGA HNSC and LUSC, which also have subtypes that express high GD2 scores ^43^. A detailed correlation analysis between the subgroups of HNSC and LUSC and BLCA subgroups would be beneficial, yet this is beyond the scope of this work.

Concerning the mechanisms related to the heterogeneous GD2 expression across tumor entities, the integration of CNA data into our analysis revealed a potential influence of *B4GALNT1* amplification on GD2 expression. *B4GALNT1* amplification was found particularly in some samples of DDLS, GBM and LUAD. In DDLS, a significant correlation was observed between samples with high-level *B4GALNT1* amplification and elevated GD2 scores. The amplification of the 12q13-15 region, which include key biomarker genes such as *MDM2*, FRS2, *CDK4*, as well as *B4GALNT1*, is a hallmark of DDLS ^53–56^. As demonstrated in Figure 8b, not all samples with *CDK4* or *MDM2* amplification exhibit co-amplification of *B4GALNT1*. This suggests the existence of distinct breaking patterns within the 12q13-15 region, with one breakpoint potentially occurring between *B4GALNT1* and *MDM2*. Given that *B4GALNT1* is a key gene of the GD2r+ variable in our model, its overexpression or amplification (or that of *ST8SIA1*) may lead to higher GD2 scores, although the implicated GD2 overexpression and the value *B4GALNT1* amplification of as surrogate biomarker of GD2 overexpression requires further validation. Moreover, our analyses in GBM and LUAD indicate that high-level *B4GALNT1* amplification does not necessarily lead to an increased GD2 score. In the GBM LGr4 subgroup, eight samples exhibited *B4GALNT1* amplification. While the difference between samples with and without this amplification was statistically significant, the overall GD2 scores were lower than those observed in DDLS. A similar pattern was observed in LUAD adenocarcinomas, where several samples showed *B4GALNT1* amplification without a corresponding significant increase in GD2 scores. Consequently, we infer that while *B4GALNT1* amplification may contribute to elevated GD2 expression in specific cancer subgroups such as DDLS, other factors likely play a more dominant role in GD2 expression regulation. Moreover, *B4GALNT1* amplification can affect GD2 expression only if the cells produce its precursor GD3 and this can be again tumor type specific.

By investigating the GD2 heterogeneity across TCGA projects and cancer subtypes, we observed a difference in the predicted expression between SCC and AD in esophageal, head and neck, endocervical, and lung cancers. An overall higher GD2 score was predicted in SCC of ESCA. SCC and AD in ESCA have specific molecular, and histopathological features ^57^. In HNSC, *B4GALNT1* was identified as a promising cell surface protein candidate for CAR-T targets based on significant expression in SCC ^58^. In the work of *Bolot et al.* ^59^, GSL expression changes in head and neck AD were reported by immunostaining and mass spectrometry. It was shown, that GM2 and GD2 were increased in these tumors compared to normal tissues.

Additionally, we observed significant differences in the predicted GD2 expression of TCGA kidney renal clear cell carcinoma across mRNA subtypes. Group 3 suggests intrasubgroup heterogeneity, where several samples exhibit high GD2 expression levels. However, the underlying drivers of GD2 expression in these subtypes remain unknown.

Currently, 23 clinical studies (basket and entity-specific) are enrolling patients for GD2- directed therapies, including CAR-T cells, monoclonal antibodies, and antibody-drug conjugates. Evaluating GD2 expression through transcriptome data can aid in accurately identifying suitable patients and provide valuable insights into interpreting clinical responses.

## 5. Conclusion

Our study provides a refined understanding of GD2 expression across various tumor entities, highlighting its biological complexity and clinical implications. We present a novel computational framework for predicting GD2-positive phenotypes using pathway-adjusted reaction activity scores based on transcriptome data. We evaluated different SVM models and achieved robust GD2 score predictions across diverse datasets and cancer types. H3K27M mutation in DMG was confirmed to be associated with elevated GD2 score and CCSK was discovered and validated as a new entity with high GD2 expression. Further, *B4GALNT1* amplification was identified as a potential GD2-promoting factor in DDLS. Additionally, we developed the GD2Viz R package to facilitate broader adoption of our method. However, GD2 scores should be interpreted with caution, as our approach relies on computational inference rather than direct GSL measurements. With our predictive GD2 score, we aim to provide insights into potential patient subgroups that could benefit from anti-GD2 therapy underscoring the relevance of GD2 as a therapeutic target in ongoing and future clinical trials.

## Code Availability

The code of our GD2Viz R package is available at https://github.com/arsenij-ust/GD2Viz. The documentation of GD2Viz can be found at https://arsenij-ust.github.io/GD2Viz/. The extended version of GD2Viz app can be found online at http://shiny.imbei.uni-mainz.de:3838/GD2Viz.

## Abbreviations

AD: Adenocarcinoma
AR: Androgen receptor
BA: Balanced accuracy
BLCA: Bladder urothelial carcinoma
BRCA: Breast cancer
CB: Cerebellum
CBTTC: Children’s Brain Tumor Tissue Consortium
CCSK: Clear cell sarcoma of the kidney
CNA: Copy number alteration
CSC: Cancer stem cells
DEA: Differential expression analysis
DDLS: Dedifferentiated liposarcoma
DIPG: Diffuse intrinsic pontine glioma
DP: Detection prevalence
DR: Detection rate
ER: Estrogen receptor
ESCA: Esophageal carcinoma
ET: Ependymoma
FDR: False discovery rate
GBM: Glioblastoma multiforme
GDC: Genomic Data Commons
GEO: Gene Expression Omnibus
GN: Ganglioneuroma
GPR: Gene-protein-reaction
GSL: Glycosphingolipid
GTEx: Genotype-Tissue Expression
HGG: High-grade glioma
HNSC: Head and neck squamous cell carcinoma
IDH: Isocitrate dehydrogenase
IHC: Immunohistochemistry
KEGG: Kyoto Encyclopedia of Genes and Genomes
LGG: Low-grade glioma
LUAD: Lung adenocarcinoma
LUSC: Lung squamous cell carcinoma
MB: Medulloblastoma
NB: Neuroblastoma
NT: Normal Tissue
RAS: Reaction Activity Score
RNA-seq: RNA sequencing
SARC: Sarcoma
SCC: Squamous cell carcinoma
SV: Support vector
SVM: Support Vector Machine
TARGET: Therapeutically Applicable Research to Generate Effective Treatments
TLC: Thin-layer chromatography
TP: Transition probability
GD2r+: GD2 promoting reactions
GD2r-: GD2 mitigating reactions

## CRediT Author Contributions

AU Conceptualization, Data curation, Formal analysis, Investigation, Methodology, Software, Visualization, Writing – original draft. FM Software, Validation, Writing – review & editing. SW Investigation, Visualization, Writing – review & editing. AW Resources, Writing – review & editing. RS Writing – review & editing. JF Resources, Writing – review & editing. CP Conceptualization, Investigation, Project administration, Supervision, Validation, Writing – original draft.

## Funding

Open Access funding enabled and organized by Projekt DEAL. We are particularly grateful for the financial support provided by the Federal Ministry of Education and Research (BMBF) in Germany, grant number: 01ZZ2322Q (PM4Onco). The work of FM was supported by the Deutsche Forschungsgemeinschaft (DFG, German Research Foundation) Projektnummer 318346496 - SFB1292/2 TP19N.

## Institutional Review Board Statement

This study was performed in agreement with the declaration of Helsinki on the use of human material for research. Ethical approval was obtained by the local ethics committee (No. 2021- 15871).

## Informed Consent Statement

In accordance with the ethics committee of Rhineland-Palatinate, written informed consent of the patient custodians was obtained for “scientific use of surplus tissue not needed for histopathological diagnosis”.

## Data Availability

This study makes use of data generated by the following St. Jude Cloud projects: St. Jude Children’s Research Hospital – Washington University Pediatric Cancer Genome Project (PCGP) and Childhood Solid Tumor Network (CSTN) ^60^, St. Jude Children’s Research Hospital Genomes for Kids Study (G4K) ^61^, St. Jude Children’s Research Hospital Real-Time Clinical Genomics (RTCG) ^62^, the Pan-Acute Lymphoblastic Leukemia Data Set of St. Jude Children’s Research Hospital (PanALL), and St. Jude Children’s Research Hospital Pediatric therapy-related myeloid neoplasm (tMN) Study ^63^.

This study makes use of the UCSC Toil RNA-seq recompute compendium ^22^.

The results published here are in part based upon data generated by the Therapeutically Applicable Research to Generate Effective Treatments (https://www.cancer.gov/ccg/research/genome-sequencing/target) initiative. The data used for this analysis are available at the Genomic Data Commons (https://portal.gdc.cancer.gov).

The Genotype-Tissue Expression (GTEx) Project was supported by the Common Fund of the Office of the Director of the National Institutes of Health, and by NCI, NHGRI, NHLBI, NIDA, NIMH, and NINDS. (https://www.gtexportal.org/)

The results published here are in part based upon data generated by the TCGA Research Network (https://www.cancer.gov/tcga).

The results published here are in part based upon data generated by the Pediatric Brain Tumor Atlas: Children’s Brain Tumor Tissue Consortium (CBTTC) (https://cbttc.org) ^30^.

The following datasets were used for validation and can be accessed via Gene Expression Omnibus (https://www.ncbi.nlm.nih.gov/geo/): GSE117446 ^31^, GSE147635 ^32^, GSE180514^16^.

## Acknowledgments

This work was supported by the computing infrastructure provided by the Core Facility Bioinformatics at the IMBEI. Further, we would like to thank Stephanie Sandor from St. Jude Cloud for her support on data availability.

## Conflicts of Interest

The authors declare no conflict of interest. The funders had no role in the design of the study; in the collection, analyses, or interpretation of data; in the writing of the manuscript; or in the decision to publish the results.

---

**Figure S1.**
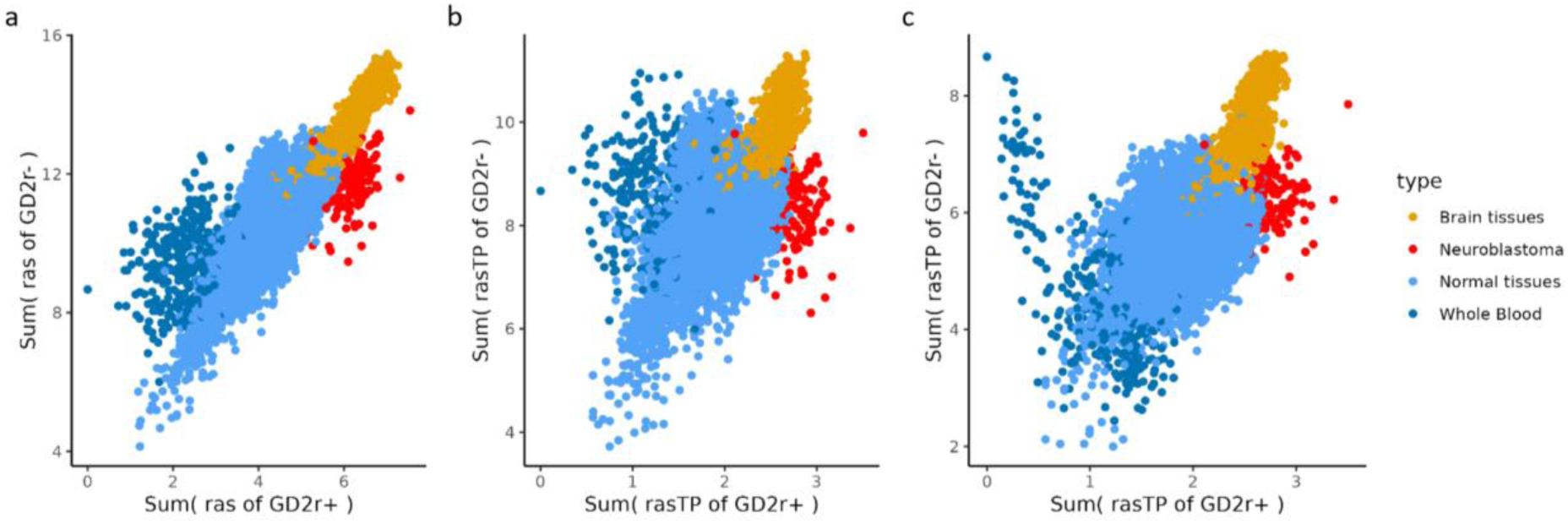
Scatterplots of the training dataset showing GD2r+ and GD2r- variables based on a) *ras*, b) *rasTP*, and c) *rasTPrec* values.

**Figure S2.**
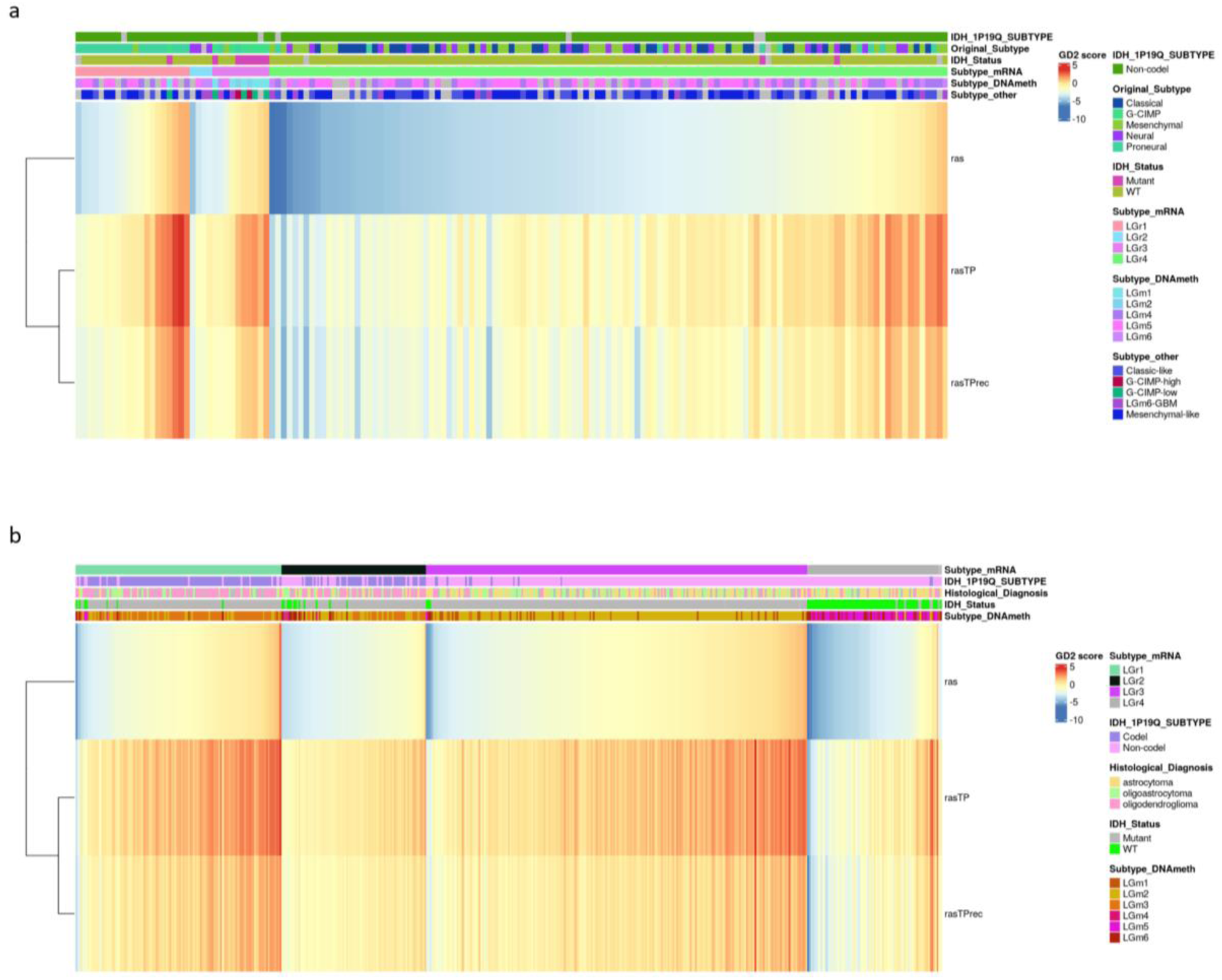
Predicted GD2 scores on *ras*, *rasTP*, and *rasTPrec* models of a) TCGA LGG and b) TCGA GBA. The heatmap is ordered based on mRNA subtype and *ras*. Sample annotation is based on *Ceccarelli et al.*.

Supplemental file 1) SVM performance results of models with different kernels. Model performance metrics for different combinations of GD2r+ and GD2r- reactions calculated on the TP-adjusted RAS values of the training dataset. #SV = Number of support vectors, Prec. = Precision, BA = Balanced Accuracy, rbfdot = Radial Basis kernel "Gaussian", polydot = Polynomial kernel, vanilladot = Linear kernel, tanhdot = Hyperbolic tangent kernel, laplacedot = Laplacian kernel, besseldot = Bessel kernel, anovadot = ANOVA RBF kernel.

Supplemental file 2) Abbreviations of cancer types and tissues used for the heatmap visualization in Figure X.

**Supplemental file 3) Statistical results for 6 datasets used for GD2 score validation.** The file contains the sample size of groups, p-values, effect sizes, and 95%CI for GD2 scores based on *ras*, *rasTP*, *rasTPrec* per dataset. The results of the Kruskal-Wallis test and the post-hoc Dunn’s test were also reported for datasets with more than two groups.

## Notes

https://github.com/arsenij-ust/GD2Viz

### Competing Interest Statement

The authors have declared no competing interest.

## References

1. Yu, R. K., Nakatani, Y. & Yanagisawa, M. The role of glycosphingolipid metabolism in the developing brain. J. Lipid Res. 50 Suppl, S440–445 (2009).

2. Yamashita, T. et al. A vital role for glycosphingolipid synthesis during development and differentiation. Proc. Natl. Acad. Sci. U. S. A. 96, 9142–9147 (1999).

3. Kasahara, K. et al. Involvement of gangliosides in glycosylphosphatidylinositol-anchored neuronal cell adhesion molecule TAG-1 signaling in lipid rafts. J. Biol. Chem. 275, 34701– 34709 (2000).

4. Sandhoff, R., Schulze, H. & Sandhoff, K. Ganglioside Metabolism in Health and Disease. Prog. Mol. Biol. Transl. Sci. 156, 1–62 (2018).

5. Sandhoff, R. & Sandhoff, K. Emerging concepts of ganglioside metabolism. FEBS Lett. 592, 3835–3864 (2018).

6. Machy, P., Mortier, E. & Birklé, S. Biology of GD2 ganglioside: implications for cancer immunotherapy. Front. Pharmacol. 14, 1249929 (2023).

7. Nazha, B., Inal, C. & Owonikoko, T. K. Disialoganglioside GD2 Expression in Solid Tumors and Role as a Target for Cancer Therapy. Front. Oncol. 10, 1000 (2020).

8. Testa, U., Castelli, G. & Pelosi, E. CAR-T Cells in the Treatment of Nervous System Tumors. Cancers 16, 2913 (2024).

9. Richards, R. M., Sotillo, E. & Majzner, R. G. CAR T Cell Therapy for Neuroblastoma. Front. Immunol. 9, 2380 (2018).

10. Das, A. K. et al. CAR T-cell therapy: a potential treatment strategy for pediatric midline gliomas. Acta Neurol. Belg. 124, 1251–1261 (2024).

11. Chan, G. C.-F. & Chan, C. M. Anti-GD2 Directed Immunotherapy for High-Risk and Metastatic Neuroblastoma. Biomolecules 12, 358 (2022).

12. Paret, C. et al. GD2 Expression in Medulloblastoma and Neuroblastoma for Personalized Immunotherapy: A Matter of Subtype. Cancers 14, 6051 (2022).

13. Vos, D. R. N., Bowman, A. P., Heeren, R. M. A., Balluff, B. & Ellis, S. R. Class- specific depletion of lipid ion signals in tissues upon formalin fixation. Int. J. Mass Spectrom. 446, 116212 (2019).

14. Panagopoulos, I., Thorsen, J., Gorunova, L., Micci, F. & Heim, S. Sequential combination of karyotyping and RNA-sequencing in the search for cancer-specific fusion genes. Int. J. Biochem. Cell Biol. 53, 462–465 (2014).

15. Rieke, D. T. et al. Feasibility and outcome of reproducible clinical interpretation of high-dimensional molecular data: a comparison of two molecular tumor boards. BMC Med. 20, 367 (2022).

16. Mabe, N. W. et al. Transition to a mesenchymal state in neuroblastoma confers resistance to anti-GD2 antibody via reduced expression of ST8SIA1. *Nat*. Cancer 3, 976– 993 (2022).

17. Ruan, S., Raj, B. K. & Lloyd, K. O. Relationship of glycosyltransferases and mRNA levels to ganglioside expression in neuroblastoma and melanoma cells. J. Neurochem. 72, 514–521 (1999).

18. Sorokin, M. et al. RNA Sequencing-Based Identification of Ganglioside GD2-Positive Cancer Phenotype. Biomedicines 8, 142 (2020).

19. Sha, Y., Han, L., Sun, B. & Zhao, Q. Identification of a Glycosyltransferase Signature for Predicting Prognosis and Immune Microenvironment in Neuroblastoma. Front. Cell Dev. Biol. 9, 769580 (2021).

20. Ustjanzew, A. et al. Unraveling the glycosphingolipid metabolism by leveraging transcriptome-weighted network analysis on neuroblastic tumors. Cancer Metab. 12, 29 (2024).

21. Goldman, M. J. et al. Visualizing and interpreting cancer genomics data via the Xena platform. Nat. Biotechnol. 38, 675–678 (2020).

22. Vivian, J. et al. Toil enables reproducible, open source, big biomedical data analyses. Nat. Biotechnol. 35, 314–316 (2017).

23. Pugh, T. J. et al. The genetic landscape of high-risk neuroblastoma. Nat. Genet. 45, 279–284 (2013).

24. GTEx Consortium. The Genotype-Tissue Expression (GTEx) project. Nat. Genet. 45, 580–585 (2013).

25. Grossman, R. L. et al. Toward a Shared Vision for Cancer Genomic Data. N. Engl. J. Med. 375, 1109–1112 (2016).

26. Carlson, M. org.Hs.eg.db. Bioconductor 10.18129/B9.BIOC.ORG.HS.EG.DB (2017).

27. Love, M. I., Huber, W. & Anders, S. Moderated estimation of fold change and dispersion for RNA-seq data with DESeq2. Genome Biol. 15, 550 (2014).

28. Colaprico, A. et al. TCGAbiolinks: an R/Bioconductor package for integrative analysis of TCGA data. Nucleic Acids Res. 44, e71 (2016).

29. Lehmann, B. D. et al. Refinement of Triple-Negative Breast Cancer Molecular Subtypes: Implications for Neoadjuvant Chemotherapy Selection. PloS One 11, e0157368 (2016).

30. Ijaz, H. et al. Pediatric high-grade glioma resources from the Children’s Brain Tumor Tissue Consortium. Neuro-Oncol. 22, 163–165 (2020).

31. Krug, B. et al. Pervasive H3K27 Acetylation Leads to ERV Expression and a Therapeutic Vulnerability in H3K27M Gliomas. Cancer Cell 35, 782–797.e8 (2019).

32. Weiss, T. et al. Schwann cell plasticity regulates neuroblastic tumor cell differentiation via epidermal growth factor-like protein 8. Nat. Commun. 12, 1624 (2021).

33. Graudenzi, A. et al. Integration of transcriptomic data and metabolic networks in cancer samples reveals highly significant prognostic power. J. Biomed. Inform. 87, 37–49 (2018).

34. Robinson, J. L. et al. An atlas of human metabolism. Sci. Signal. 13, eaaz1482 (2020).

35. Goksuluk, D. et al. MLSeq: Machine learning interface for RNA-sequencing data. Comput. Methods Programs Biomed. 175, 223–231 (2019).

36. Benjamini, Y. & Hochberg, Y. Controlling the False Discovery Rate: A Practical and Powerful Approach to Multiple Testing. J. R. Stat. Soc. Ser. B Stat. Methodol. 57, 289– 300 (1995).

37. Galassi, L. et al. Naxitamab Activity in Neuroblastoma Cells Is Enhanced by Nanofenretinide and Nanospermidine. Pharmaceutics 15, 648 (2023).

38. Mount, C. W. et al. Potent antitumor efficacy of anti-GD2 CAR T cells in H3-K27M+ diffuse midline gliomas. Nat. Med. 24, 572–579 (2018).

39. El Malki, K., et al. Glucosylceramide Synthase Inhibitors Induce Ceramide Accumulation and Sensitize H3K27 Mutant Diffuse Midline Glioma to Irradiation. Int. J. Mol. Sci. 24, 9905 (2023).

40. Wei, X., Li, S. & Wang, Y. Expression of GD2 and GD3 in peripheral neuroblastic tumors. Indian J. Pathol. Microbiol. (2024) doi:10.4103/ijpm.ijpm_618_23.

41. Ceccarelli, M. et al. Molecular Profiling Reveals Biologically Discrete Subsets and Pathways of Progression in Diffuse Glioma. Cell 164, 550–563 (2016).

42. Cancer Genome Atlas Research Network. Comprehensive molecular characterization of clear cell renal cell carcinoma. Nature 499, 43–49 (2013).

43. Cancer Genome Atlas Research Network. Comprehensive molecular characterization of urothelial bladder carcinoma. Nature 507, 315–322 (2014).

44. Benedetti, D. J. et al. Treatment and outcomes of clear cell sarcoma of the kidney: A report from the Children’s Oncology Group studies AREN0321 and AREN03B2. Cancer 130, 2361–2371 (2024).

45. Cutcliffe, C. et al. Clear cell sarcoma of the kidney: up-regulation of neural markers with activation of the sonic hedgehog and Akt pathways. Clin. Cancer Res. Off. J. Am. Assoc. Cancer Res. 11, 7986–7994 (2005).

46. Yu, R. K., Bieberich, E., Xia, T. & Zeng, G. Regulation of ganglioside biosynthesis in the nervous system. J. Lipid Res. 45, 783–793 (2004).

47. Yu, R. K., Nakatani, Y. & Yanagisawa, M. The role of glycosphingolipid metabolism in the developing brain. J. Lipid Res. 50 Suppl, S440–445 (2009).

48. Straathof, K. et al. Antitumor activity without on-target off-tumor toxicity of GD2- chimeric antigen receptor T cells in patients with neuroblastoma. Sci. Transl. Med. 12, eabd6169 (2020).

49. Majzner, R. G., Weber, E. W., Lynn, R. C., Xu, P. & Mackall, C. L. Neurotoxicity Associated with a High-Affinity GD2 CAR-Letter. Cancer Immunol. Res. 6, 494–495 (2018).

50. Jennemann, R., Rodden, A., Bauer, B. L., Mennel, H. D. & Wiegandt, H. Glycosphingolipids of human gliomas. Cancer Res. 50, 7444–7449 (1990).

51. Battula, V. L. et al. Ganglioside GD2 identifies breast cancer stem cells and promotes tumorigenesis. J. Clin. Invest. 122, 2066–2078 (2012).

52. Ly, S. et al. Anti-GD2 antibody dinutuximab inhibits triple-negative breast tumor growth by targeting GD2+ breast cancer stem-like cells. J. Immunother. Cancer 9, e001197 (2021).

53. Dei Tos, A. P. Liposarcoma: new entities and evolving concepts. Ann. Diagn. Pathol. 4, 252–266 (2000).

54. Conyers, R., Young, S. & Thomas, D. M. Liposarcoma: molecular genetics and therapeutics. Sarcoma 2011, 483154 (2011).

55. Nishio, J., Nakayama, S., Nabeshima, K. & Yamamoto, T. Biology and Management of Dedifferentiated Liposarcoma: State of the Art and Perspectives. J. Clin. Med. 10, 3230 (2021).

56. Jing, W. et al. Expression of FRS2 in atypical lipomatous tumor/well-differentiated liposarcoma and dedifferentiated liposarcoma: an immunohistochemical analysis of 182 cases with genetic data. Diagn. Pathol. 16, 96 (2021).

57. Cancer Genome Atlas Research Network et al. Integrated genomic characterization of oesophageal carcinoma. Nature 541, 169–175 (2017).

58. Park, Y. P. et al. CD70 as a target for chimeric antigen receptor T cells in head and neck squamous cell carcinoma. Oral Oncol. 78, 145–150 (2018).

59. Bolot, G. et al. Analysis of glycosphingolipids of human head and neck carcinomas with comparison to normal tissue. Biochem. Mol. Biol. Int. 46, 125–135 (1998).

60. Downing, J. R. et al. The Pediatric Cancer Genome Project. Nat. Genet. 44, 619–622 (2012).

61. Newman, S. et al. Genomes for Kids: The Scope of Pathogenic Mutations in Pediatric Cancer Revealed by Comprehensive DNA and RNA Sequencing. Cancer Discov. 11, 3008–3027 (2021).

62. Rusch, M. et al. Clinical cancer genomic profiling by three-platform sequencing of whole genome, whole exome and transcriptome. Nat. Commun. 9, 3962 (2018).

63. Schwartz, J. R. et al. The acquisition of molecular drivers in pediatric therapy-related myeloid neoplasms. Nat. Commun. 12, 985 (2021).

